# A network medicine approach to elucidate mechanisms underlying menopause-induced knee osteoarthritis

**DOI:** 10.1101/2023.03.02.530756

**Authors:** Gabrielle Gilmer, Hirotaka Iijima, Natalie Jackson, Zachary Hettinger, Allison C. Bean, Juliana Bergmann, Christopher Evans, Rebecca C. Thurston, Fabrisia Ambrosio

**Affiliations:** Medical Scientist Training Program, School of Medicine, University of Pittsburgh, Pittsburgh, PA, USA; Cellular and Molecular Pathology Graduate Program, University of Pittsburgh, Pittsburgh, Pittsburgh, PA, USA; Discovery Center for Musculoskeletal Recovery, Schoen Adams Research Institute at Spaulding, Boston, MA, USA; Department of Physical Medicine & Rehabilitation, Harvard Medical School, Boston, MA, USA; Department of Geriatrics, University of Pittsburgh, Pittsburgh, PA, USA; Institute for Advanced Research, Nagoya University, Nagoya University, Nagoya, Japan; Department of Bioengineering, University of Pittsburgh, Pittsburgh, PA, USA; Department of Geriatric Medicine, University of Pittsburgh. Pittsburgh, PA, USA; Department of Physical Medicine and Rehabilitation, University of Pittsburgh, Pittsburgh, PA, USA; McGowan Institute for Regenerative Medicine, University of Pittsburgh, Pittsburgh, PA, USA; Department of Biological Sciences in the Dietrich School of Arts & Sciences, University of Pittsburgh, Pittsburgh, PA; Department of Physical Medicine & Rehabilitation, Mayo Clinic, Rochester, MN, USA; Department of Psychiatry, University of Pittsburgh School of Medicine, Pittsburgh, PA, USA

## Abstract

Post-menopausal women present with the highest incidence and morbidity of knee osteoarthritis (KOA), but no disease-modifying therapies are available. This treatment gap may be driven by the absence of menopause in preclinical studies, as rodents do not naturally maintain a menopausal phenotype. Here, we employed a chemically-induced menopause model to map the trajectory of KOA at the tissue and proteome levels and test therapeutics *in silico*. Middle-aged female mice were randomized to sesame oil (non-menopause) or 4-vinycyclohexene diepoxide (menopause) injections. Following comprehensive validation of our model, knees were collected across perimenopause and menopause for histology, and cartilage samples were micro-dissected for mass spectrometry proteomics. Menopause mice displayed aggravated cartilage degeneration and synovitis relative to non-menopause mice. An unbiased pathway analysis revealed progesterone as a predominant driver of pathological signaling cascades within the cartilage proteome. Network medicine-based analyses suggested that menopause induction amplifies chondrocyte senescence, actin cytoskeleton-based stress, and extracellular matrix disassembly. We then used *in silico* drug testing to evaluate how restoration of sex hormones impacted the cartilage network. The greatest restoration was observed with combined estradiol/progesterone treatment (i.e., hormone therapy), although *in silico* treatment with a senolytic drug also partially recovered the cartilage proteome. Taken together, our findings using a translatable female aging model demonstrate that menopausal aging induces progressive cartilage degeneration and amplifies age-related synovitis. These changes may be driven by a previously unappreciated role of progesterone loss and menopause-induced cellular senescence. Lastly, *in silico* treatment suggests an estradiol/progesterone cocktail or senolytics may attenuate menopause-induced cartilage pathology.

**One Sentence Summary:** Menopause induces cartilage degradation, senescence, and extracellular matrix disassembly, while hormone therapy restores the cartilage proteome.

## INTRODUCTION

In 1843, Søren Kierkegaard reflected, “*What a curse to be a woman! And yet the very worst curse… is, in fact, not to understand [woman].”*(*1*) Two hundred years later, the medical and biological community still lack a fundamental understanding of female physiology. Many factors contribute to this persistent gap, including a lack of sex-specific pharmacological targets (*2*), limited inclusion of women in clinical trials (*3, 4*), and underfunding of research for diseases disproportionately affecting women.(*5*) Another major, yet far less appreciated, contributing factor to our incomplete understanding of females is that pre-clinical models all-too-often fail to recapitulate diseases as they present in women. This translational disconnect is particularly problematic in the context of aging female biology. At 9-12 months of age, female rodents experience ‘estropause’ which is the cessation of estrus cyclicity, similar to the cessation of menstrual cycles seen in humans with the onset of menopause.(*6*) However, within months, 75% of rodents spontaneously rejuvenate their ovarian follicles.(*7*) In these cases, sex hormones and estrus cyclicity mirror pre-menopause, despite the inability to breed.(*8*) Given women spend approximately one third of their life in menopause (*9*), the absence of menopause in animal models obstructs translatability of preclinical work for half the population.

One consequence of this insufficiency of female aging animal models is a gap in the ability to treat diseases associated with menopause, such as knee osteoarthritis (KOA).(*10*) Although KOA is twice as likely to present in post-menopausal women than in men (*11, 12*), a recent systematic review showed that approximately 75% of murine studies of KOA only included males.(*13*) Of the few studies that did include females and menopause to study KOA, the predominant menopause model used was ovariectomy (OVX).(*7, 14*) However, the fidelity with which OVX recapitulates natural menopause is questionable. Specifically, OVX results in abrupt session of ovarian function, lacks a perimenopausal period (i.e., the gradual transition from regular menstrual cycles to a menopausal state lasting 7-12 years in humans (*15*)) and disrupts all ovarian sex hormones, including ones that do not normally change in menopause, such as testosterone.(*7*) Additionally, OVX studies for KOA have largely employed young animals(*14*), despite clinical evidence clearly showing that young women who have oophorectomies and older women undergoing natural menopause represent distinct populations with distinct clinical needs.(*16*) As such, OVX is unlikely to provide the much-needed biological insights related to natural menopause. The vast majority of clinical trials aimed to treat KOA have failed in phase I or II(*17*), and the scarcity of both females and natural menopause models in preclinical KOA studies is likely contributing to this shortfall.

With this background in mind, the purpose of this study was to evaluate the effects of the natural menopause transition on the trajectory of KOA pathogenesis in a murine model. We started by validating a chemically-induced menopause model in middle-aged female mice. Since KOA is considered a disease of the whole joint,(*18*) we thoroughly documented histological changes in cartilage, synovium, and subchondral bone across perimenopause and menopause.

We then performed mass spectrometry proteomics on micro-dissected cartilage to track changes in proteomic pathways over the menopausal transition and applied network-medicine approaches to our mass spectrometry data with the goal of modeling treatment modalities *in silico*. Of note, throughout the text we refer to humans with uteruses using the terms ‘woman’ and ‘women’. We recognize that these terms contain gender information, but for the purpose of our discussion, these terms indicate humans that undergo natural menopause.

## RESULTS

### VCD model of menopause in middle-aged female mice recapitulates a natural menopause phenotype without off target effects

We first validated the VCD model of menopause in middle-aged female C57BL/6 mice using previously established protocols.(*8*) Unlike OVX, VCD retains intact ovaries and induces a perimenopausal transition.(*8*) Enrolled mice were randomized to receive intraperitoneal (IP) injections of either VCD in sesame oil or sesame oil only for 10 consecutive days (**Figure 1A**).(*8*) We used 14-16 month old mice since this age corresponds to 47-52 years old in humans, which is when most women begin the menopausal transition.(*19*)

**Figure 1:**
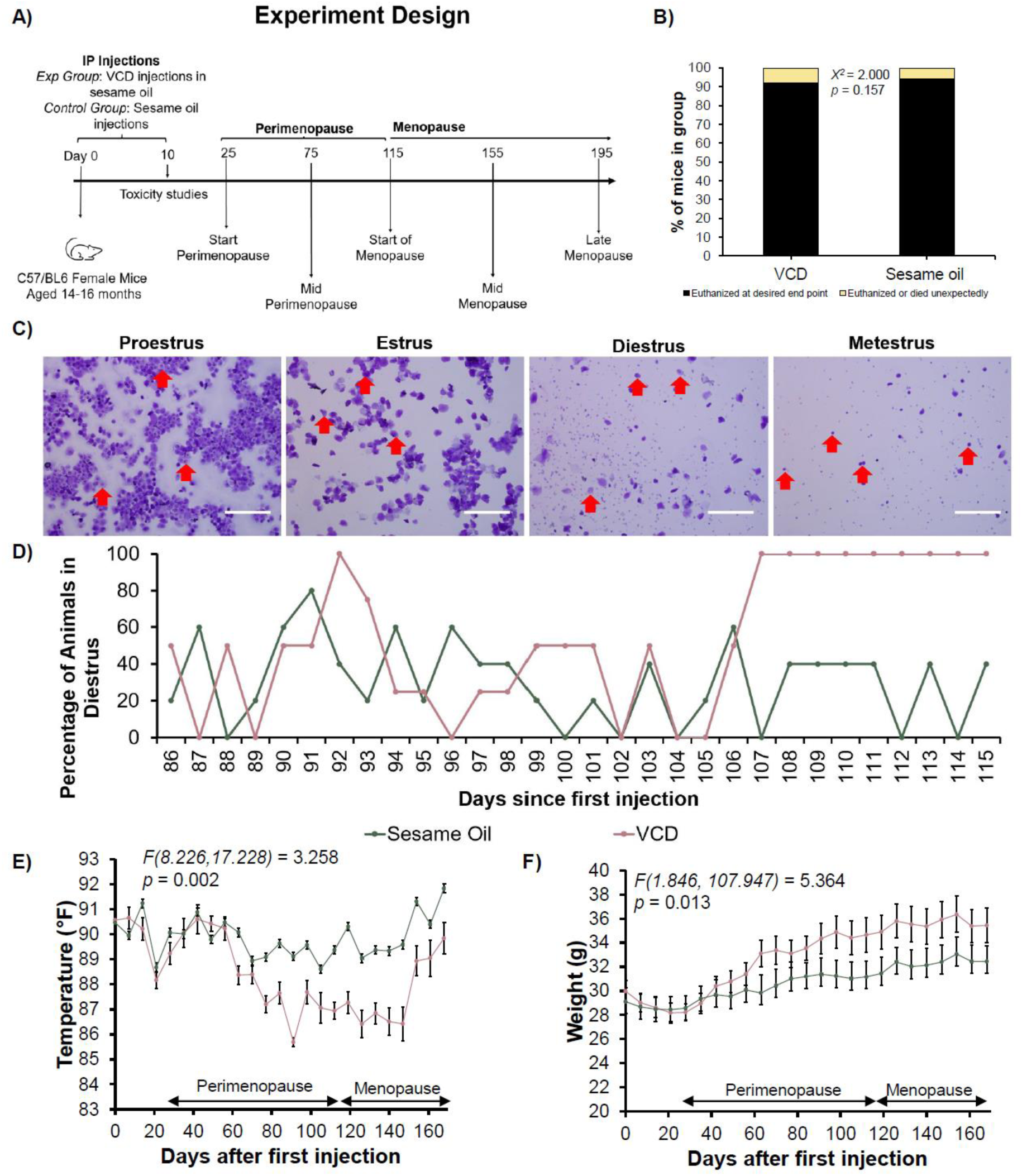
4-Vinylcyclohexene diepoxide (VCD) model of menopause recapitulates vaginal cytology, temperature, and weight changes seen in menopause with no off-target effects. A) Experimental design, intraperitoneal injection (IP) schedule, euthanasia schedule, and sample size per time point. At day 10, n=3/group. At day 25, n=5/group. At day 75, n=7 for VCD and 5 for sesame oil. At day 115, n = 7/group. At day 155, n =6 in VCD group and n=5 in sesame oil group. At day 195, n=6/group. B) Unexpected deaths per group. There were no differences in toxicity or mortality based on injection type (Supplementary File 1). C) Representative images of each phase of the estrus cycle. In proestrus, red arrows indicate nucleated epithelial cells. In estrus, red arrows represent cornified epithelial cells. In diestrus, red arrows represent leukocytes and nucleated epithelial cells. In metestrus, red arrows indicate leukocytes. Scale bar = 100 μm. D) The percentage of sesame oil and VCD injected mice in the diestrus phase of the estrus cycle. Menopause onset was defined as ten days in a row of diestrus. VCD injected animals become menopausal 115 days after the first injection, while sesame oil animals continue to have regular estrus cycles. E) Weekly rectal temperature recordings (mean±95%CI) from VCD and sesame oil injected mice. VCD mice showed a drop in body temperature over the course of perimenopause that recovered at the onset of menopause. F) Weekly weight recordings (mean±95%CI) from VCD and sesame oil injected mice. While both groups gradually gained weight with time, the VCD mice gained weight at a faster rate than the sesame oil mice. All reagents/materials needed to perform these experiments are listed in Table S6, and all statistical tests are listed in Table S7.

Within the VCD and sesame oil injected animals, 8% and 6% of mice, respectively, died unexpectedly or were euthanized owing to adverse events (**Figure 1B**). Internal organs from all mice in these studies were evaluated by a pathologist for assessment of off-target effects from VCD or sesame oil. A blinded pathologist concluded that lesions (e.g., mild multifocal lymphoplasmacytic, histiocytic, and neutrophilic infiltrates in the peritoneal fat) in both the VCD and sesame oil animals were most likely due to repeated IP injections (**Supplementary File 1**). These findings are in line with previous toxicology reports in young mice(*20*) but contrasts those documented in adult rats.(*21*) Specifically, while no signs of toxicity beyond the ovaries were noted in young mice,(*8*) a previous report indicated that 100% of adult rats developed peritonitis from injections of VCD.(*21*)

VCD animals presented with cycle irregularity (defined as estrus cycles 6-7 days or longer(*7*)) 25±3 days after the first injection and became menopausal 115±2 days after the first injection (defined as 10 consecutive days of the diestrus phase of the estrus cycle) (*7, 8, 22*) (**Figure 1C-D**). Conversely, sesame oil animals continued to have regular estrus cycles (cycle length: 5.23±0.64 days.(*23*) Having validated that VCD mice present with a menopausal phenotype while the sesame oil mice do not, from this point forward we refer to VCD mice as “menopause group” and the sesame oil mice as “non-menopause group”. Also, of note, for the remainder of our experiments, we defined the “start of perimenopause” as the point when menopause mice presented with irregular cycles (day 25), and the “start of menopause” as the point when the menopause mice presented with 10 consecutive days in diestrus (day 115). “Mid- perimenopause” and “mid-menopause” were defined as the middle points between the start and end/late of perimenopause and menopause, respectively. “Late menopause” was defined as the animals reaching 21 months of age, which corresponds to approximately 60 years in humans and represents that age when notable changes in response to hormone therapy for treatment of menopause-related symptoms in humans are observed (*10*) (**Figure 1A**).

To further characterize how well the VCD model mimics natural menopause, rectal temperatures and body weights were documented weekly. The menopause group had a significant drop in body temperature during perimenopause that returned to the level of the non-menopause group with the onset of menopause (**Figure 1E**). Neff et al. reported that body temperature is lower in women undergoing perimenopause,(*24*) and irregularities in body temperature maintenance are highly associated with vasomotor symptoms.(*25, 26*) Interestingly, in previous studies using OVX, authors reported an *increase* in body temperature following OVX compared to sham.(*27, 28*) It is unclear if this discrepancy in temperature change is due to differences in physiological adaptations to these models or experimental differences, such as measuring temperature at different times of day.(*29*) The menopause group also had a steady rise in body weight that outpaced the non-menopause group (**Figure 1F**). Specifically, the menopause group gained weight at a rate of 0.34 g per 9 days or 1.13% weight gain per 9 days relative to the average starting weight of 30 g (∼9 mouse days ∼= 1 human year.(*30*)) Similarly, women undergoing menopause gain weight at approximately 1.5 pounds per year or 0.88% weight gain per year relative to an average starting weight of 170 pounds.(*31, 32*) Menopause is associated with increases in peri-abdominal and visceral fat(*33*), and our finding is consistent with these clinical observations.

### Menopause-induction compromised cartilage and synovium health

To ensure that VCD and sesame oil were not directly affecting joint tissues, an initial cohort of joints were collected immediately after completion of injections (day 11). We detected no differences in histological score of cartilage, synovium, or subchondral bone at this early time point (**Figure S1**). We next collected knees from menopause and non-menopause groups across the perimenopause and menopause transition (**Figure 1A**). When examining changes in cartilage histology across time, the menopause group presented with progressively increasing Osteoarthritis Research Society International (OARSI) scores from mid-perimenopause to late menopause. A higher OARSI score indicates more severe cartilage degeneration.(*34*) In contrast, there were no differences in OARSI scores in the non-menopause group at any of the timepoints evaluated. Relative to the non-menopause group, the menopause group had significantly higher OARSI scores at the start of menopause (*U* = 5.545, *p* = 0.019), mid-menopause (*U* = 7.500, *p* = 0.006), and late menopause (*U* = 7.500, *p* = 0.006) (**Figure 2A-B, Figure S2A**). Since we observed more weight gain in the menopause animals, we also repeated this analysis with weight normalized scores, which did not change the conclusion (**Figure S2B**). These findings suggest that menopausal aging induces progressive cartilage degeneration, while non-menopausal aging does not and are in line with previous work.(*35*) Additionally, the OARSI scores in the menopause, but not the non-menopause, group are in line with radiographic tracking of cartilage degeneration in women(*36–39*), suggesting the menopause group presents with a representative phenotype of KOA in women.

**Figure 2:**
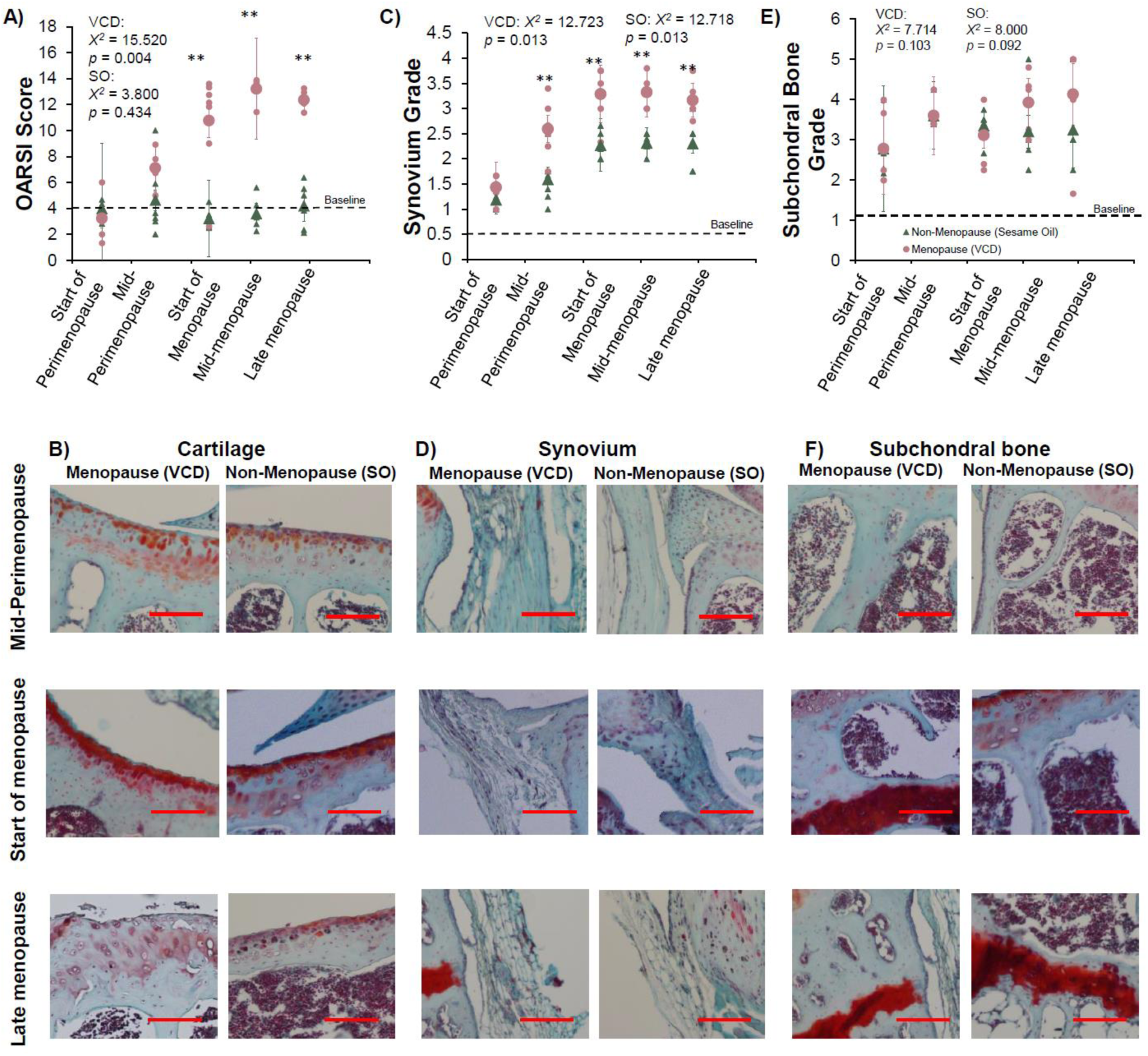
Menopausal aging results in progressive cartilage and synovium pathology compared to non-menopausal aging. A) Osteoarthritis Research International Society (OARSI) scoring of cartilage degeneration. The non-menopause (sesame oil injected, SO) group demonstrated no changes in cartilage degeneration across time, while the menopause (4-vinylcyclohexene diepoxide, VCD) group demonstrated significantly increased cartilage degeneration across time and compared to the non-menopause group. B) Representative images of cartilage between groups at mid- perimenopause, start of menopause, and late menopause. C) Grade of synovium pathology. Both the non-menopause and menopause groups presented with progressive synovium pathology across time, but the menopause group was more severe than the non-menopause group. D) Representative images of the synovium between groups at mid-perimenopause, start of menopause, and late menopause. E) Grade of subchondral bone pathology. There were no differences between the menopause and non-menopause groups nor any differences across time. F) Representative images of the subchondral bone between groups at mid-perimenopause, start of menopause, and late menopause. Dashed lines indicate baseline degeneration observed in middle-aged mice immediately after injections (**Figure S1**). Representative images from start of perimenopause and mid-menopause are available in **Figure S3**. ** - indicates significant difference between groups (menopause vs non-menopause) at specific time point. The statistical values listed on the graphs are for the interaction of time. All reagents/materials needed to perform these experiments are listed in **Table S6**, and all statistical tests are listed in **Table S7**. Data are presented as mean ± standard deviation. Scale bar = 100 μm.

Both menopause and non-menopause groups presented with significant increases in synovium grade over time (**Figure 2C-D**), indicative of progressive thickening and/or inflammation.(*40*) However, the menopause group had significantly worse synovitis than the non-menopause group at mid-perimenopause (*U* = 5.810, *p* = 0.016), start of menopause (*U* = 6.403, *p* = 0.011), mid-menopause (*U* = 7.604, *p* = 0.006), and late menopause (*U* = 5.791, *p* = 0.016) (**Figure 2C-D, Figure S2C**). As before, we repeated this quantification with weight normalized scores and saw no change in overall conclusion (**Figure S2D**). These findings suggest that aging propagates synovium pathology, and menopause amplifies this effect.

Lastly, we quantified changes in subchondral bone pathology. We observed no effects of time or group on raw or weight normalized scores (**Figure 2E-F, Figure S2E-S2F**). Previous studies using both VCD(*20, 41*) and OVX(*42*) models have reported changes in subchondral bone pathology with induction of menopause. However, different methodologies to quantify subchondral bone pathology were used (e.g., bone mineral density via DEXA), and the animals were young at the time of menopause onset. Additionally, given that our power analysis was based on cartilage outcomes, it is possible we were underpowered to evaluate differences in bone health.

### Menopause did not induce changes in frailty, grip strength, or wheel running

Menopause models in animals are associated with behavioral changes(*43*), and KOA is affected by activity level(*44–46*), with “too much” (e.g., high level athletes) or “too little” (e.g., sedentary lifestyle) physical activity increasing risk of KOA. Thus, one possible explanation for the aforementioned histological findings is that menopause induces changes in activity level, ultimately leading to detrimental effects to the cartilage and synovium. To test this possibility, we evaluated whether menopause results in changes to activity level, as determined by grip strength, frailty, and wheel running behavior. The menopause and non-menopause groups had similar all-paw and hind-paw grip strengths, frailty indexes, and wheel running behaviors before IP injections. However, forepaw strength was slightly higher in the menopause group compared to the non-menopause group (**Figure S3**). Evaluation of these behaviors again at the start of menopause revealed no differences between groups in any of the parameters (**Figure 3**). This finding suggests that the histological changes presented here are unlikely to be due to activity level changes.

**Figure 3:**
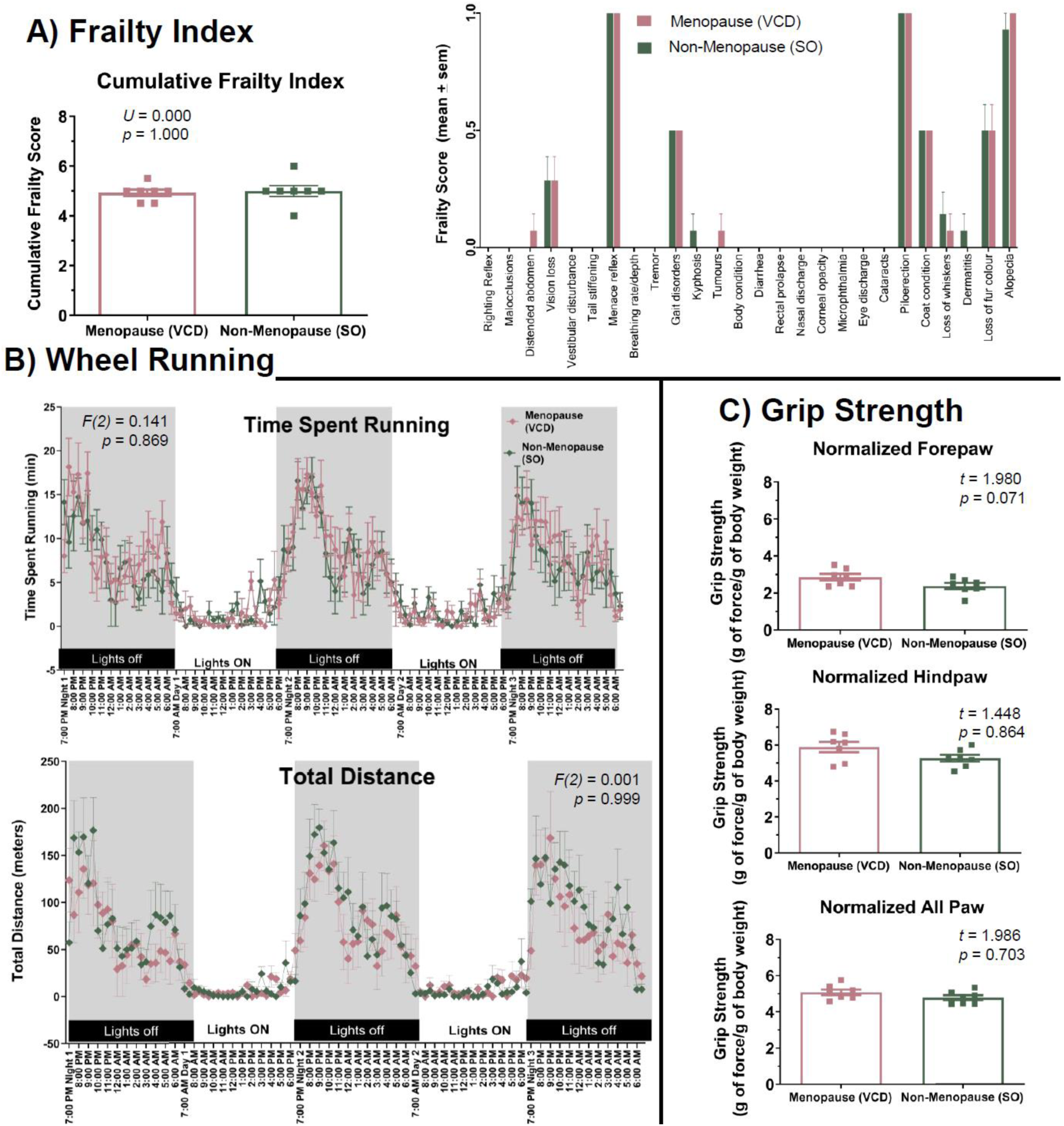
Menopause-onset did not induce changes to frailty, wheel running, or grip strength at the start of menopause. All behavioral tasks were evaluated at the start of menopause, all groups had n =7, and data are presented as mean ± standard error of the mean. All reagents/materials needed to perform these experiments are listed in **Table S6**, and all statistical tests are listed in **Table S7**. There were no differences between any of the behavioral assessments between groups. A) Cumulative frailty index and individual scores of the different contributing components. Example frailty index scoring sheet is available in **Supplementary File 2**. B) Time spent and distance run during the wheel running assay. C) Forepaw, hindpaw, and all-paw grip strength normalized to body weight.

### The proteomic pathway “oocyte meiosis” is associated with menopause induction in cartilage

To augment our mechanistic understanding of menopausal effects on KOA, we turned our attention to hypothesis-generating techniques. Standard techniques currently exist for isolating cartilage from murine knees,(*47*) but technical limitations hamper reproducible extraction of the synovium from mouse joints.(*48*) As such, we focused on further characterizing menopausal effects on the cartilage using mass spectrometry proteomics. We collected cartilage samples from menopause and non-menopause groups at mid-perimenopause (i.e., the time point right before the statistical changes), start of menopause, and late menopause in order to better understand the long-term trajectory of KOA (**Figure 4A**).

**Figure 4:**
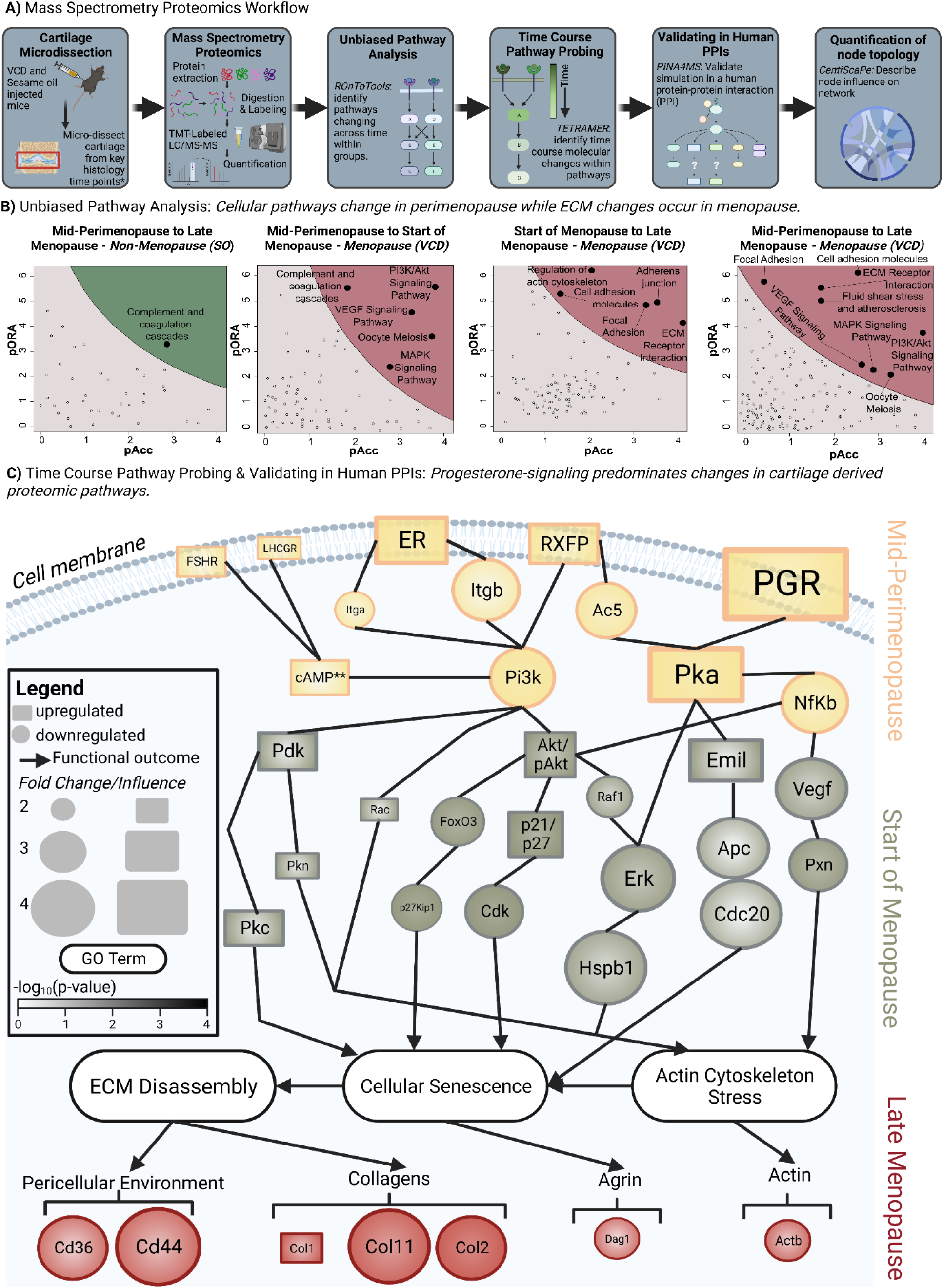
Menopause induction disrupts progesterone driven proteomic pathways and leads to cellular senescence, actin cytoskeleton stress, and extracellular matrix (ECM) disassembly. A) Mass spectrometry proteomics workflow. Briefly, cartilage was micro-dissected from the knees of menopause (4-vinylcyclohexene diepoxide, VCD) and non-menopause (sesame oil, SO) mice at time points determined from Figure 3. LC/MS-MS mass spectrometry proteomics with TMT-labeling was performed. An unbiased Kyto Encyclopedia Genes and Genomics (KEGG) Pathways Analysis, time course simulation (Temporal TRAnscriptional regulation ModellER (TETRAMER)), validation of protein-protein interactions (PPI) in human networks (Protein Interaction Network Analysis For Multiple Sets (PINA4MS)) (Figure S4B), and quantification of node topology (CentiScaPe) (Figure S4C-S4E) were performed. B) Results from unbiased KEGG Pathway Analysis using ROnToTools. pORA represents the Boolean value of over- representation p-value, and pAcc represents the Boolean value of total bootstrap permutations accumulation. The remaining results from the non-menopause group are in Figure S4A. C) Summary of time course pathway probing. Each node represents a summary of proteins (e.g., Itgb represents Itgb1, Itgb3, Itgb5, Itgb6, Itgb7, Itgb4, Itgb8), and lines represent a PPI. Full list of summarized proteins is available in Table S3. The top 10 Gene ontology (GO) terms with associated p-values are listed in Table S4. Shape indicates directionality of change, color indicates time point in which change occurred (defined as greater than 2-fold change), and size indicates fold change (menopause to non-menopause). Figure S4B displays the network representing nodes that were directly detected in the mass spectrometry data (versus implied from the model and downstream effects) and lines symbolize PPI (FigureS4B for representation of each database).

TMT-labeled mass spectrometry proteomics identified 5,343 unique proteins from 768,579 M2 spectra. We first performed an unbiased Kyoto Encyclopedia of Genes and Genomes (KEGG) Pathway Analysis using ROnToTools.(*49*) In the non-menopause group, the only pathway changing across time was *Complement and coagulation cascades* (**Figure 4B, S4A**), consistent with a previous report(*35*). Conversely, in the menopause group, changes were progressive. From mid-perimenopause to the start of menopause, changes in cell-based pathways predominated, including *PI3K/Akt Signaling* and *MAPK Signaling*, while from the start of menopause to late menopause, changes occurred in ECM-associated pathways, including *Focal Adhesion* and *Cell Adhesion Molecules* (**Figure 4B**). This suggests that in menopause-induced KOA, dysregulation within the resident cells in cartilage, chondrocytes, proceeds—and may facilitate—ECM dysfunction, which has been supported by other studies.(*50*) Another possible explanation for this finding is that menopause induces a shift in cartilage composition from a proportionally more “cell based” composition (i.e., a higher cell to ECM ratio) to a proportionally more “ECM based” composition (i.e., a lower cell to ECM ratio).(*51*) As such, there is a need for more proteomic analyses with specific attention paid to fractionating cell and ECM based components as well as other -omics techniques, such as RNA-sequencing, to interrogate these observations.

Interestingly, of the pathways identified, the only one clearly regulated by sex-hormones was *Oocyte meiosis* (see **Figure 4B**). Additionally, *Oocyte meiosis* was noted to be changing from mid-perimenopause to the start of menopause (i.e., before the statistically significant change in cartilage histology and before the changes in ECM based pathways). The *Oocyte meiosis* KEGG Pathway is composed of progesterone-regulated MAPK signaling and protein ubiquitination. Given the early changes in this pathway in our analyses and the early drop of progesterone in perimenopause,(*52*) this finding implies a novel role of progesterone signaling in mediating menopause-related changes to the cartilage proteome.

### A network medicine approach reveals progesterone and estradiol signaling modulate cellular senescence, actin cytoskeleton stress, and ECM disassembly in cartilage

We next integrated the aforementioned pathways together into a single “network” to identify how individual pathway elements change with time.(*53*) Most molecules exert their functions through exchanges with other molecules within the same cell, with neighboring cells, or with distant cells. A network medicine approach maps these interactions to elucidate how abnormal signaling impacts pathology.(*53*) Specifically, we used Temporal TRAnscriptional regulation ModellER (TETRAMER) to simulate the pathways identified in our ROnToTool analysis.(*54*) Given our focus on menopause, we also integrated pathways known to be regulated by sex hormones (e.g., estradiol, progesterone, relaxin). We then validated our network in human protein-protein interaction (PPI) networks using the Protein Interaction Network Analysis For Multiple Sets (PINA4MS)(*55, 56*) (see **Methods** for details on simulations). A complete list of included pathways, input materials, and an abbreviation key are available in **Tables S1-S3**.

At mid-perimenopause, the menopause group demonstrated alterations in cellular signaling regulated by integrins (Itga, Itgb), nuclear factor-κB (NFκB), and phosphoinositide 3- kinases (PI3K) relative to the non-menopause group (**Figure 4C**). At the start of menopause, our simulations suggested changes in focal adhesion (Pxn), actin reorganization (Pkn, Hspb1, Rac), regulation of chondrogenesis (Pkc), and chondrocyte cell cycle/proliferation (Erk, Cdk, p27Kip1) occur in the menopause group relative to the non-menopause group. Lastly, in late menopause, the consequences of these maladapted signaling cascades were observed with increased expression of type I collagen (Col1) at the expense of decreased expression of pericellular environment stabilizers (Cd44, Cd36), agrin (Dag1), actin (Actb), and type II and XI collagens (Col2, Col11) in the menopause group relative to the non-menopause group (**Figure 4C, S4B**). These findings were summarized using g:Profiler(*57, 58*) and can be categorized into three predominant gene ontology (GO) biological processes: cellular senescence, actin cytoskeleton stress, and ECM disassembly (**Figure 4C**, Top 10 GO Terms and p-values listed in **Table S4**).

Our simulations also implied that sex hormone signaling, in rank order, was most significantly mediated by (1) progesterone (PGR), (2) estradiol (ER), (3) relaxin (RXFP), (4) follicle stimulating hormone (FSH, FSHR), and (5) luteinizing hormone (LH, LHCGR). These effects were quantified by effect size, betweenness (i.e., capacity of a protein to communicate with distant proteins(*59*)), and centroid (i.e., probability of a protein to be functionally capable of organizing downstream proteins(*59*)) (**Figure S4C-E**). Although many studies have implied that loss of estradiol(*42, 60–62*) and heightened FSH levels(*63–65*) may drive KOA, these findings implicate a previously unrecognized role of disrupted progesterone, relaxin, and LH signaling in mediating cartilage homeostasis.

### *In silico* treatment with combined progesterone and estradiol partially restores the cartilage proteome

To better define how these sex hormones mediate menopause-driven cartilage degeneration and to “treat” our network, we performed *in silico* simulations using NeDRex. NeDRex is an integrated package in Cytoscape for evaluating potential drug repurposing opportunities.(*66*) Specifically, we simulated how administration of Cetrorelix (LH-receptor antagonist(*67*)), Suramin (FSH-receptor antagonist(*68*)), Serelaxin (relaxin supplement (*69*)), Estradiol, Micronized Progesterone, Progesterone/Estradiol (combined Levonorgestrel + estradiol treatment), and Raloxifene (Selective Estrogen Receptor Modulator) altered network topology. Based on our network medicine analyses (**Figure 4C**), we hypothesized that Micronized Progesterone would have the strongest effects, followed by Estradiol, Raloxifene, Serelaxin, Suramin, and Cetrorelix.

We quantified the effects of each drug on the network by first comparing the fold change in individual proteins making up the network. We defined “low impact” drugs as having an average fold change of < 1.4, “moderate impact” drugs as having an average fold change of 1.4 < *x* < 1.7, and “high impact” drugs as having an average fold change > 1.7 relative to the untreated network. Additionally, we quantified how each treatment modulated the overall network “cohesiveness” in CentiScaPe(*59*) by calculating network closeness, radiality, and diameter.

Networks with high closeness and radiality are more likely to organize and “lead” protein signaling, while low closeness and radiality implies an open system that connects driving systems to outcomes.(*59*) Diameter is a measure of the overall “easiness” of proteins to communicate and influence their downstream signaling cascades.(*59*) Lastly, we quantified changes in biological processes following drug treatment with g:Profiler as was performed in our initial network analyses.(*57*)

As expected, Cetrorelix and Suramin had low impacts on proteomic signaling cascades (**Figures S5A-S5B**). Despite the relatively strong effect of relaxin and estradiol in our above simulations **(Figure 4C**), Serelaxin, Estradiol, and Raloxifene also had low impact on restoring the cartilage proteome (**Figures S5C, S6A-S6B**). Micronized Progesterone treatment had moderate impacts (**Figure S6C**), while Progesterone/Estradiol had high impacts on the cartilage proteome network (**Figure 5A**). When looking at effects on overall network topology relative to no treatment, we saw that Progesterone/Estradiol consistently led to a more cohesive network, as evidenced by heightened closeness, radiality, and diameter (**Figures 5B-D**). Lastly, when looking at biological processes changing as a result of drug treatment, we saw only one change in the top 10 GO biological process with *in silico* treatment of Cetrorelix and Suramin, and only 3-4 changes with *in silico* treatment of Serelaxin, Estradiol, Raloxifene, and Micronized Progesterone (**Table S4**). Conversely, treatment with combined Progesterone/Estradiol changed 6 of the top pathways (**Table S4**). Specifically, the terms “cellular senescence”, “actin cytoskeleton stress”, and “ECM disassembly” were no longer in the top 10. Instead, these terms were replaced with “negative regulation of cellular senescence”, “actin cytoskeleton organization”, and “gene expression involved in ECM organization” following treatment.

**Figure 5:**
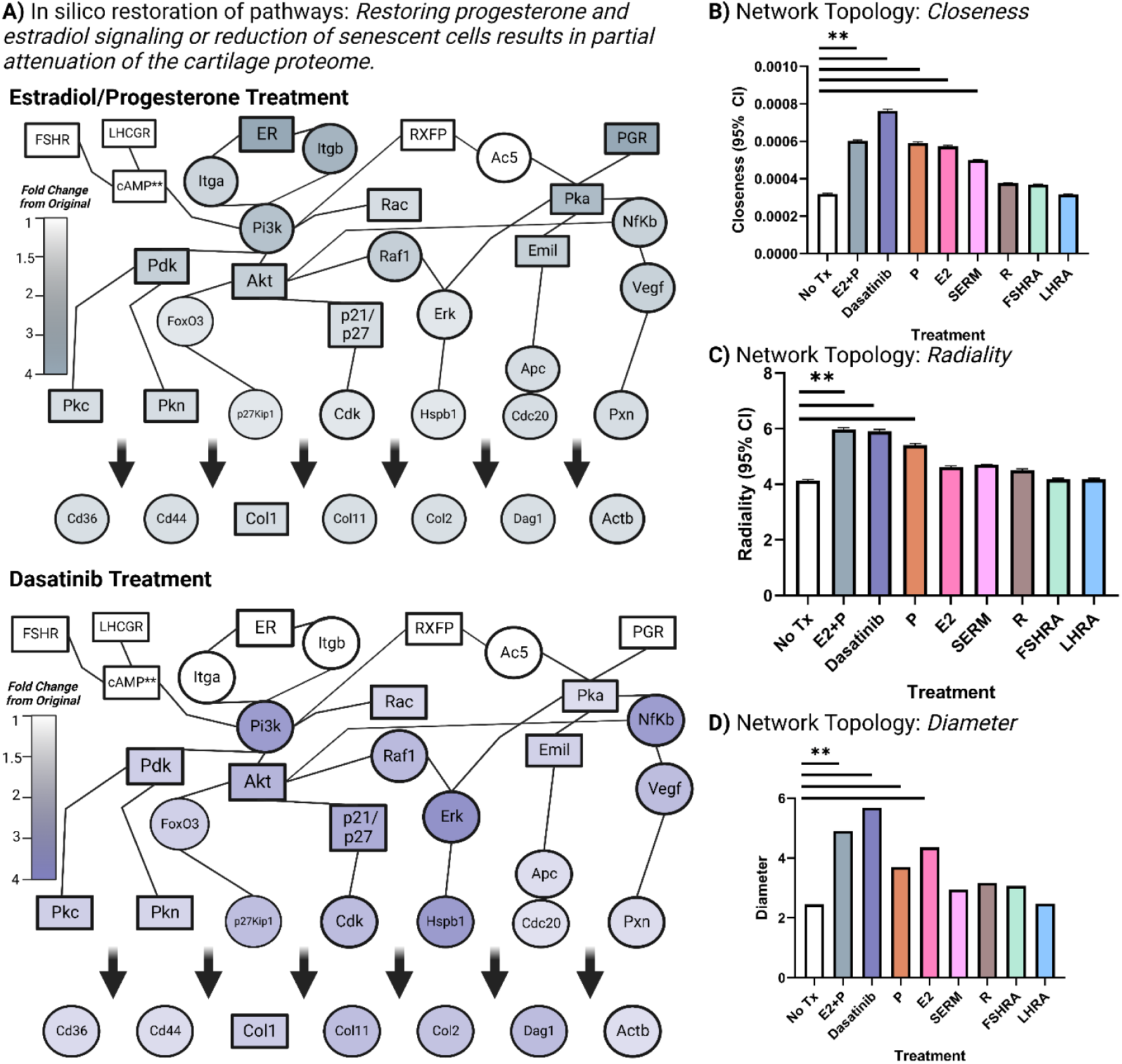
*In silico* treatment with combined Progesterone/Estradiol or a senolytic drug results in partial restoration of the cartilage proteome. A) Results of *in silico* drug stimulation using combined Progesterone/Estradiol (E2+P) treatment and Dasatinib (senolytic drug) treatment. Fold change is relative to the untreated network shown in Figure 4C. Other drug simulations are reported in Figure S5. B) Quantification of closeness (i.e., a signaling network with high closeness is more likely to organize functional units or modules, whereas a signaling network with low closeness will behave more like an open cluster of proteins connecting different regulatory modules) with no treatment (No Tx), E2+P, Dasatinib, Micronized Progesterone (P), Estradiol (E2), Serelaxin (R), Suramin (FSHRA), and Cetrorelix (LHRA) treatments. C) Quantification of radiality (i.e., a signaling network with high radiality is more likely organizing functional units or modules, whereas a signaling network with low radiality will behave more like an open system) with No Tx, E2+P, Dasatinib, P, E2, R, FSHRA, and LHRA treatments. D) Quantification of diameter (the overall easiness of the proteins to communicate and influence their downstream function) with No Tx, E2+P, Dasatinib, P, E2, R, FSHRA, and LHRA treatments. ** means p < 0.05. All statistical tests are listed in Table S7. Created in BioRender.

The culmination of our findings suggests that restoring isolated sex-hormone signaling is unlikely to rejuvenate the cartilage proteome to a non-menopausal state. Indeed, when comparing the difference between the isolated estradiol and progesterone treated networks together versus the combined Progesterone/Estradiol treated network, we saw that effects were synergistic, not just additive, in nature, specifically along MAPK signaling (**Figure S6D**). Interestingly, combined Progesterone/Estradiol treatment offered some benefits to the cartilage proteome. This finding implies that menopause-induced KOA may be driven, in part, by dysregulation in estradiol-progesterone-cross talk and that restoration of this interaction may be a promising therapeutic target.

### *In silico* treatment with a senolytic drug partially restores the cartilage proteome

Given the changes in cellular senescence seen in our GO analysis, we also simulated how administration of Dasatinib, a senolytic drug, affected our network.(*70*) When looking at effects on overall network topology relative to no treatment, we saw that Dasatinib drove a more cohesive network, as quantified by heightened closeness, radiality, and diameter (**Figures 5B-D**). When considering biological processes, treatment with Dasatinib changed eight of the top pathways (**Table S4**). Specifically, Dasatinib treatmens offered benefits to the cartilage proteome that were similar in effect on individual proteins, biological process, and network topology as the combined Progesterone/Estradiol treatment. This finding supports that regulating senescence may be a promising modality for the treatment of OA in menopausal females.

## DISCUSSION

In this paper, we outlined changes associated with menopause-induced KOA from the molecular to whole organism level. First, we showed that the VCD model in middle-aged female mice closely mimics the phenotype observed in menopausal women, as determined by loss of estrus cyclicity, thermal dysregulation, and gradual weight gain. We then demonstrated that menopause propagates cartilage degeneration and worsens age-related synovium pathology.

However, menopause onset did not result in changes to frailty, grip strength, or wheel running, suggesting that menopause-induced KOA was not driven by menopause-associated activity changes. Early changes in progesterone-signaling impeded proteomic pathways associated with cellular senescence and ECM maintenance. Lastly, *in silico* “treatment” of combined Progesterone/Estradiol or a senolytic drug resulted in partial restoration of the cartilage proteome, which may translate to improved cartilage quality and attenuation of menopause-induced KOA. (**Figure 6**)

**Figure 6:**
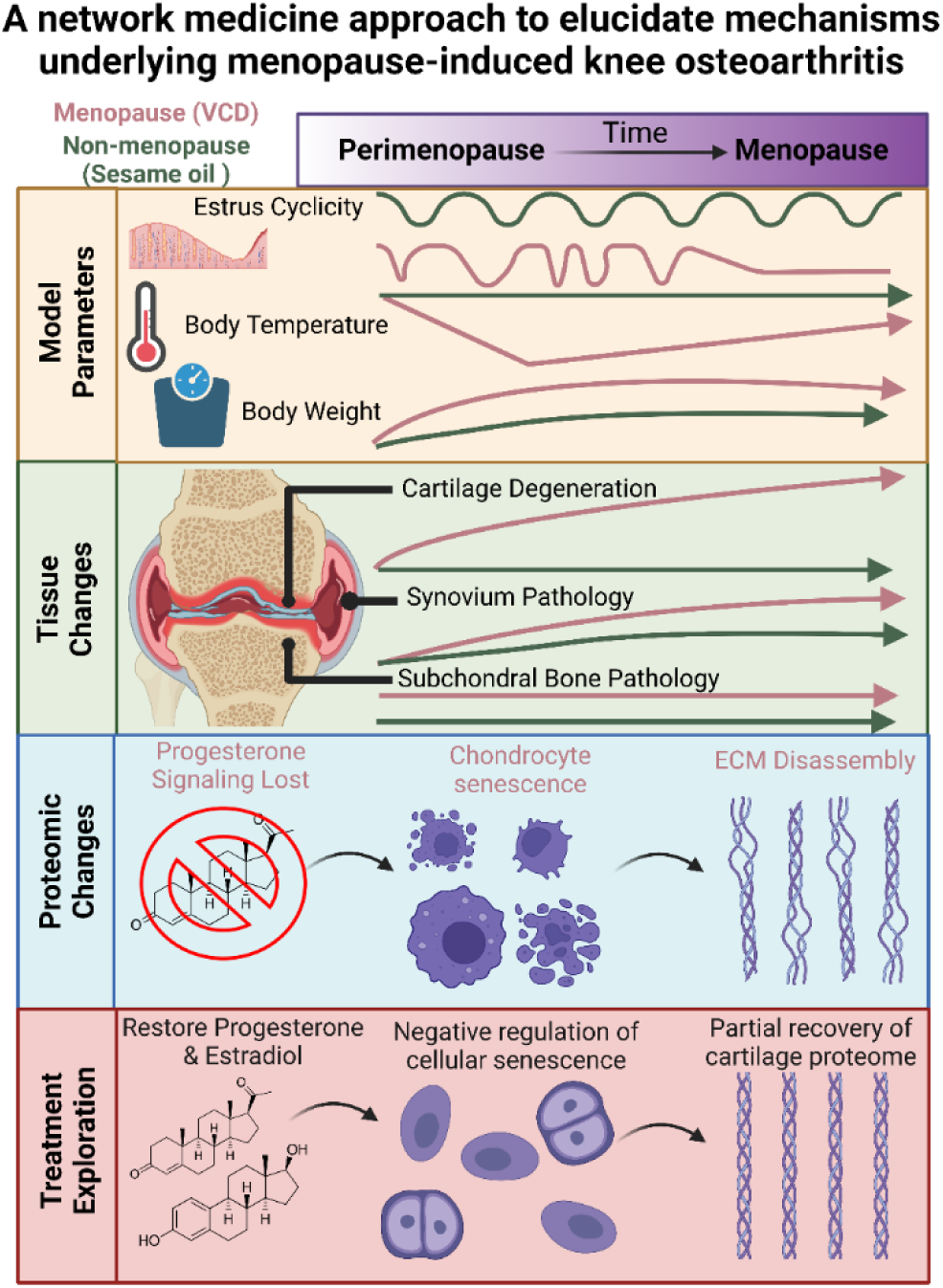
Graphical abstract for *A network medicine approach to elucidate mechanisms underlying menopause-induced knee osteoarthritis*. *Model Parameters:* We characterized the 4-vinylcyclohexene diepoxide (VCD) model of menopause in middle-aged female mice. Sesame oil (non-menopause) animals continued to have estrus cycles, while VCD (menopause) animals demonstrated cycle irregularity and ultimately cycle cessation. Body temperature decreased in perimenopause and recovered during menopause in the menopause group while the non-menopause group remained constant. Weight gain in the menopause group outpaced that of the non-menopause group. *Tissue Changes:* The menopause group demonstrated progressive cartilage degeneration starting in mid-perimenopause, while the non-menopause group did not. Synovium pathology in the menopause group outpaced that seen in the non-menopause group. No subchondral bone pathology was detected in either group. *Proteomic Changes:* Of the sex-hormones known to change in menopause, loss of progesterone signaling was predicted to have the largest effects on proteomic networks within cartilage. Disrupted cellular signaling pathways were noted to change in mid-perimenopause, while proteomic pathways related to the extracellular matrix (ECM) were found to change late in menopause. Summarized gene ontology (GO) terms revealed cellular senescence and ECM disassembly were prominently affected biological processes. *Treatment Exploration: In silico* treatment with combined progesterone and estradiol partially restored the cartilage proteome and alleviated pathological biological processes. Created in BioRender.

Assessment of non-cartilage tissues is becoming standardized practice within osteoarthritis research, and in humans, there is strong evidence that the synovium propagates cartilage related pathologies.(*71*) We observed in these studies a statistical difference in the synovium that preceded cartilage alterations. However, the cartilage and synovium association was only observed in the menopause mice. That is, whereas synovium pathology was detected in both the menopause and non-menopause groups, cartilage integrity was only disrupted in menopause animals. This interesting observation advocates for the study of the synovium-cartilage axis in the context of menopause to better understand the interactions driving the phenotypes observed.

When looking more closely at the ECM proteins that changed with menopause, we observed type XI collagen was most affected. Type IX and XI collagens maintain the spacing and structural distribution between type II collagen fibers, which is the predominate collagen in cartilage.(*72*) The consequences of losing this structural regulation has been studied in humans, with single nucleotide polymorphisms of Col11a1 being associated with KOA.(*73*) In some cancers, intervertebral disc disease, and retinal pathologies, changes in type XI collagen mediate changes in stiffness and drive maladaptive mechanotransductive mechanisms in the resident cell populations.(*74–76*) Thus, one hypothesis generated from our findings is that menopause-induced chondrocyte senescence leads to pathologic turnover of type XI collagen. This maladaptive environment could further disturb chondrocyte homeostasis, thus creating a feedforward loop of worsening cartilage degeneration, as has been suggested in other KOA studies.(*77*) With this potential implication, future work into these specific mechanisms is warranted.

At the level of the cartilage, PI3K/Akt signaling regulates ECM anabolism, chondrocyte proliferation and apoptosis, autophagic flux, and inflammation.(*78–80*) In our recent work, we showed that PI3K/Akt signaling was disrupted in male, but not non-menopausal female, mice with aging.(*35*) PI3K/Akt signaling is known to be regulated by estradiol.(*81*) Thus, our findings support the role of estradiol regulating PI3K/Akt signaling within menopause-induced cartilage degeneration. Relaxin, a peptide hormone similar in structure to insulin, is also known to regulate fibrosis via PI3K/Akt signaling.(*82*) Of note, relaxin regulates ECM stiffness and mediates collagen turnover within cartilage, as has been shown using pregnancy models (*83–87*). Our findings further support this role; however, given most of the prior work has been performed in the context of pregnancy, there is a need for targeted studies aimed at better understanding the role of relaxin in menopause-induced KOA.

In breast cancer, it is well-known that progesterone-driven MAPK signaling mediates disease progression(*88–91*), though the role of MAPK in the context of KOA is less appreciated. Our findings highlight a novel role of this signaling cascade in menopause-induced KOA. There is some clinical evidence that progesterone, especially when combined with estradiol, may induce protective effects against OA.(*92, 93*) However, to our knowledge, this is the first study to dissect underlying mechanisms and to highlight the prominence of progesterone signaling in cartilage degeneration. Indeed, in our unbiased pathway analysis, *Oocyte meiosis*, which is predominantly regulated by progesterone, was the only direct sex-hormone pathway to continuously change throughout the evaluated time course. Similarly, *in silico* simulations revealed that progesterone treatment in isolation demonstrated more beneficial effects to the cartilage proteome than estradiol or SERM treatment alone. Thus, our findings spotlight the need for more mechanistic studies aimed at elucidating the role of progesterone as well as sex hormone interactions in mediating menopause-induced KOA.

There is a large body of evidence suggesting that chondrocyte senescence is a driver of KOA(*94–101*), though the underlying mechanisms are not well-established. Some studies have suggested chondrocyte senescence is driven by Forkhead transcription factors(*94*), secretory factors from other joint tissues,(*95*) accumulation of reactive oxygen species and mitochondrial dysfunction,(*96*) genomic and epigenomic damage,(*97*) and other pathologic signaling cascades. (*98, 99, 101*) Our findings add to this body of literature by showing an association between menopause and the loss of progesterone signaling with senescence. This latter association was further supported by our *in silico* simulations showing Dasatinib partially alleviates the cartilage proteome in a manner similar to combined progesterone/estradiol treatment. Dasatinib acts via regulation of Src and modulation of PI3K(*102*) MAPK(*103*) and Nfkb(*104*), three pathways seen in our network. In breast cancer, progesterone is known to regulate the Src/p21ras/Erk pathway via cross-talk with estradiol(*105*), and our findings suggest a similar response in the context of menopause-driven cartilage degeneration. Although these findings are based on simulations, they represent a valuable opportunity for mechanistic exploration and treatment potential.

Although this study adds to a growing body of literature aimed at understanding menopause-induced KOA, there are limitations. There is a large body of evidence that histopathological outcomes do not necessarily correlate with pain, which is the predominant complaint of clinical KOA.(*106*) It is unclear how this disconnect affects our study outcomes and translation of our findings. Another limitation of this work is that the mechanistic insights and “treatment” modalities were assessed *in silico*. Specifically, *in silico* treatments modeled how a given intervention affected the network, but this approach does not take into account the nuance of timing of treatment initiation,(*14*) systemic factors, or the downstream impact on tissue morphology and symptom presentation.(*107*) While network medicine approaches are robust and have shown great success in identifying molecular insights(*53*), there is a need to validate these pathway findings and treatments *in vivo* and *in vitro*.

Two hundred plus years have passed since Kierkegaard aptly pointed out the curse caused by not understanding women, and the road ahead to enmesh our animal models to the diseases that present in post-menopausal women remains long. We anticipate that these studies will provide a road map for further mechanistic studies into the impacts of menopause on KOA and towards the development of disease-modifying therapeutics.

## MATERIALS AND METHODS

All reagents and equipment used in our studies are outlined in **Table S6**. *Note*: Supplementary tables and data will be made available upon publication of this work. Please contact the corresponding author for more information before then.

### Animal handling & safety

All animals enrolled in our experiments were 14–16-month-old female C57/BL6 mice obtained from the National Institute on Aging (NIA) Rodent Colony. Mice were housed in cages with a temperature (22°-23°C), humidity (55-65%), and light (12-hour light/dark cycle) controlled environment. Domes were placed in cages for enrichment, food and water access was ad libitum, and 3-5 mice were housed per cage. All animal experiments outlined here were approved by the University of Pittsburgh’s Institutional Animal Care and Use Committee.

If severe reactions or pain were noticed during any of the below procedures, including lethargy, poor grooming, 20% weight loss, or hunching, a veterinarian was consulted immediately to determine an appropriate plan. If any mouse was found dead unexpectedly or required euthanasia prior to the planned endpoint, they were excluded from the knee and behavioral analyses.

Internal organs (heart, lungs, liver, spleen, kidneys, perineal fat, stomach, large intestine, small intestine, pancreas) of all animals, including from those animals with unexpected endpoints, were fixed in 10% formalin, prepared in paraffin blocks, sectioned with a microtome, stained using Hematoxylin and Eosin, and assessed for toxicity by a blinded pathologist.

### 4-Vinylcyclohexene Diepoxide (VCD) Model of Menopause

Mice were acclimated to the environment for at least two days prior to any handling. Since there has been documentation of differences in behavior based on the sex of the investigator(*108*), mice were only handled by females. All handling and intraperitoneal (IP) injections took place between 4:30-6:30 am EST by two investigators (GG and NJ). To acclimate the mice to the mechanical restrainer, the investigators handled the mice for five days by placing them in a mechanical restrainer and then placing them back in their cage prior to any injections beginning. The mechanical restrainer is a clear, tube-like structure with a narrow hole on the bottom to allow for IP injections, and a “door close” that allows the investigator to secure the tail and access the rectum and vagina. On the last day of handling, mice were randomized via coin flip to receive IP injections of either VCD in sesame oil or just sesame oil. Once a group had half the possible animals via coin flip, the remaining animals were assigned to the other group (e.g., for a total N = 16, if after 13 coin flips, the VCD group = 8, and the sesame oil group = 5, the remaining 3 animals were assigned to the sesame oil group). All animals were housed in the same room in the animal facility, and cage location within that room was randomized via a random number generator.

VCD was given at a dose of 160 mg/kg/day suspended in sesame oil via IP injection using a well-established protocol.(*8*) The total volume of all injections (both sesame oil and VCD) ranged from 70-80 μL depending on the mouse’s weight. On the morning of each set of injections, a coin was flipped to determine which group would be injected first (VCD or sesame oil). Syringes and needles were prepped using sterile technique and one needle/syringe set was used per mouse. Each mouse was placed in the mechanical restrainer. Injection placement alternated between the right and left lower quadrant to minimize irritation with consecutive days of injections. The injection site was defined as the level of the hip/at the level of the second set of nipples and was cleaned with an alcohol pad. One investigator lifted the restrainer such that the mouse’s tail was above her head. The other investigator inserted the needle, going no deeper than half of the needle inserted, bevel up, at a 30-40° angle. Mice were temporarily placed in a tall bucket for monitoring of behavior and bleeding. If bleeding was noticed, the area was cleaned with an alcohol wipe and the animal was checked again later in the day. These steps were repeated for 10 consecutive days.

### Rectal Temperature, Weight Monitoring, & Vaginal Cytology and Lavage

Rectal temperatures and weights were monitored weekly. Tape was used to mark a 5 cm depth on the rectal thermometer to ensure consistent and repeatable measurements. To measure temperature, each mouse was placed in the mechanical restrainer on the same day each week at 4:30 am EST. After securing the mouse, a rectal thermometer was inserted into the mouse’s rectum and held steady until the temperature readout remained constant for five seconds. The investigator then removed and cleaned the probe with an alcohol wipe prior to taking a reading on a different animal. Mice were then placed individually on a laboratory scale for weight measurement. Weights were recorded once the mouse was standing fully on the scale.

Vaginal lavage and cytology were performed based on an established protocol.(*22*) For vaginal lavage, double distilled water (ddH2O) was autoclaved and stored at room temperature until collections. All vaginal lavage collections took place between 4:30-6:30 am EST and were performed daily until the mice injected with VCD became menopausal. Approximately 200 μL of ddH2O was drawn up into two different plastic transfer pipets. Each mouse was individually placed within the mechanical restrainer. Once the mouse was secure and ceased urination and defecation, a pipet was used to wash the genital area. After cleaning, the other ddH2O filled pipet was placed at the vaginal canal, with attention paid to not disturb the opening. The pipet bulb was gently depressed to dispense approximately 50-100 μL at the opening of the vaginal canal, while the vagina spontaneously aspirated the fluid. The pipet bulb was then released, pulling the ddH2O back into the pipet. This step was repeated 15-20 times. After the final collection of the ddH2O back into the pipet, the fluid was then dispersed onto a clean, glass slide. Three slides were collected per day per mouse. Each mouse was then removed from the restrainer and placed back into their cage.

Slides were allowed to dry at room temperature for at least one day prior to staining. Crystal violet stain was prepared by mixing 0.1% of crystal violet powder in ddH2O. Slides were stained in crystal violet in a Tissue-Tek slide staining rack for one minute. Slides were then washed twice in ddH2O for 1 min each. Excess water and stain were removed from the edges using a Kimwipe.

Slides were immediately imaged at 10X using a light microscope by a blinded investigator (NJ). Three images were taken per slide with attention paid to capture the representative ratios of cornified squamous epithelial cells, leukocytes, and/or nucleated epithelial cells. A blinded reviewer (ACB) assessed the phase of the estrus cycle using the ratio of cells types present as outlined in this previous protocol.(*22*)

### Histological preparation of mouse knee joints and sectioning

All mice were euthanized between 4:30-6:30 am EST and kept on ice until knees were collected. Left knee joints were prepared for histological assessment as outlined in previous studies.(*35*) Briefly, the skin at the ankle was removed to expose the entire leg. Muscles, ligaments, tendons, and fat were carefully removed using dissecting scissors, with attention paid to neither cut the bone nor penetrate the synovium. Of note, to not disturb the synovium, the patella and patella tendon were not removed. After excess tissue was removed, the knee joint was separated from the leg by cutting at the mid-femur and mid-tibia. Knees were fixed in 2 mL of 4% paraformaldehyde (PFA) on a shaker at 4°C overnight. Knees were then washed with phosphate buffered solution (PBS) for 5 minutes, three times and transferred to 10 mL of decalcification solution. Decalcification solution was changed every three days, and knees were incubated for a total of ten days (i.e., start incubation on day 1, change solution on day 4 and day 7, and remove knees from solution on day 10). Knees were again washed with PBS three times for 5 minutes. Dehydration of the knees began by incubating the knees in 25% and 50% ethanol for 2 hours each at room temperature, followed by incubating knees in 70% ethanol overnight at 4°C. The following day, knees were incubated in 95% and 100% ethanol for 2 hours each at room temperature and then placed into glass tubes for incubation in Xylene for 1 hour at room temperature. Xylene was removed and fresh Xylene was added for another hour of incubation.

Xylene was again removed, and knees were incubated in liquid paraffin at 60°C for 2 hours. Knees were then cut along the medial collateral ligament (MCL) by pinning the tibia and femur with needles on an Xacto cutting board such that the MCL was facing up. Using a microtome blade, one investigator carefully cut along the MCL to generate an anterior and posterior plane of the knee. Knee halves were placed into a fresh container of liquid paraffin and incubated overnight at 60℃.

Samples were embedded by placing the cut plane of the knee face down into a paraffin base mold. An embedding cassette was placed over the sample, and the cassette/mold was filled with liquid paraffin wax. Samples were refrigerated overnight, after which samples were stored at room temperature until use. Sections were made 6 μm thick, with four sections per slide and 15 slides were collected per sample. After a section was generated, it was placed in a hot water bath for 2-3 minutes to prevent any wrinkles. Sections were placed on slides using a small, wooden rod. Slides were allowed to dry at least overnight prior to staining.

### Safranin-O and Fast Green Staining

Slides were incubated in a 60℃ oven overnight in a slide holder. Paraffin was cleared by placing slides in Histoclear twice for 5 minutes. Slides were then successively dipped in 100%, 95%, 70%, and 50% ethanol for 2 minutes each, followed by washing for 1 min in ddH2O. Slides were stained in hematoxylin for 1 minute, followed by a 10 min wash of running tap water, with water flowing to the back side of the slides. The investigator dipped the slides in 1% acid ethanol (1 mL HCl in 400 mL of 70% ethanol) and then dipped the slides in 0.01% fast green for 2 minutes. Slides were dipped in 1% acetic acid for 10 seconds and incubated in 0.5% Safranin-O for 1 hour. An investigator rehydrated the slides in 95% and 100% ethanol two times for 5 minutes each. Lastly, slides were placed in Histoclear twice for 10 minutes each. Slides were carefully dried using a Kimwipe and mounted using Limonese-Mount.

### Histological scoring of cartilage, synovium, and subchondral bone

Images of the stained medial tibial cartilage were taken at 10X using a standard bright field microscope by a blinded investigator (NJ). This area was chosen since it is the second most affected area in humans(*109*) and mouse models historically have recapitulated this severity.(*110*) Of note, the most affected area in humans is the patellofemoral joint, which tends to not present with severe OA, likely due to different joint mechanics between biped versus quadruped, and is also difficult to visualize and study in mice due to size. The Osteoarthritis Research Society International (OARSI) Histopathology Initiative scoring system was used to evaluate the grade (severity) and stage (diffuseness) of cartilage lesions, the grade of the synovium, and the grade of the subchondral bone.(*111*) Seven different areas of the medial knee joint (one image per slide) were evaluated, and the scores were averaged across these different areas. A trained, blinded examiner (JB) evaluated the histological sections (intraclass correlation coefficient, 95% confidence interval: 0.878, 0.799 - 0.915 with another reviewer (ACB))

### Behavioral Assays

The cumulative frailty index was assessed using methods outlined previously(*112*) and an example scoring sheet is provided in **Supplemental File 3**. Briefly, mice were individually assessed by a blinded behavioral expert who is trained in assessment for frailty evaluation. The investigator scored the absence, presence, and/or severity of 27 different characteristics associated with frailty. Each item was given a 0, 0.5, or 1 based on the severity, and items were summed to give the cumulative frailty index. Maximum score is a 27, indicating highest level of frailty.

The protocol for assessment of wheel running was derived from a previous study, and the investigator performing the assessment was blinded to the groups.(*112*) Given limitations in cage size, the food hopper was removed and placed on the floor, and mice were housed individually. The running wheel is a 15.5-cm diameter plastic plate that is mounted to an electronic base. On the underside of the wheel, there is a small magnet that is tracked by the electronic base. The electronic base is battery powered, monitors wheel revolutions continuously, and transmits data to a system acquisition every 30 seconds. Total time and total distance run on the wheel were recorded and analyzed.

Grip strength was evaluated using previously established protocols, and the investigator performing the assessment was blinded to the groups.(*112*) Briefly, the investigator allowed the mouse to grab onto the wire mesh with either the forepaws, hind paws, or all four paws. The mouse was then pulled by either the tail (for forepaw) or neck scruff (for hindpaw and all paw) until it released its grip. The wire mesh pad was connected to a force transducer which measured the maximum force during each iteration. Three consecutive trials of each condition were recorded, normalized to body weight, and averaged.

### LC/MS-MS mass spectrometry-based proteomics

Using previously established protocols(*35, 113*), cartilage was microdissected from the right knee of female C57/BL6 mice who received either sesame oil or VCD injections. Samples were collected at Mid-Perimenopause (day 70), Start of Menopause (day 115), and Late Menopause (day 195). Briefly, the skin was carefully pulled back from the ankle to expose the entire leg.

Muscles, fat, ligaments, and tendons were carefully removed from the knee joint and surrounding area. Once the joint line was visible, microdissection scissors were used to separate the femoral condyles from the tibial plateau. Both surfaces were then placed in PBS under a dissecting microscope. The surface was visualized, and microdissection scissors and tweezers were used to remove the menisci, ligaments, fat, and synovium. After the surface was clear, scissors were used to scrape cartilage from the surface of the joint. Collected cartilage was washed in PBS and then placed in a collection vial. This was repeated on the tibial plateau. Samples were stored at - 80℃ until all samples were collected. At this point, samples were lyophilized and shipped to the Washington University at St. Louis Proteomics and Mass Spectrometry Program (PMSP).

Cartilage samples were prepared for mass spectrometry proteomics using methods previously described.(*35*) Briefly, cartilage samples were homogenized, sonicated, and clarified. Lysates was reduced with TCEP and alkylated with 500mM 2-Chloroacetamide. Lysates were digested in LysC, diluted, and again digested overnight in Trypsin. Digestions were stopped by acidifying, and desalting was performed on C18 spin columns (cat. 89870, Fisher Scientific) according to manufacturer instructions. Peptides were lyophilized and resuspended in 0.1% formic acid. Peptide counts were quantified with the Pierce Quantitative Colorimetric Peptide Assay.

15 µg of peptides from each sample were labeled with TMTpro reagents dissolved in anhydrous acetonitrile for 1 hour at room temperature. Three runs were performed for each time point, with both groups present. Each fraction was resuspended in 20 µl 0.2% formic acid.

Liquid chromatography-mass spectrometry (LC-MS) analysis of peptide fractions was carried out on an EASY-nLC 1000 coupled to an Orbitrap Eclipse Tribrid mass spectrometer (Thermo Fisher Scientific). Each fraction was loaded onto an Aurora 25cm x 75µm ID, 1.6µm C18 reversed phase column (Ion Opticks, Parkville, Victoria, Australia) and separated over 136 min at a flow rate of 350 nL/min with the following gradient: 2–6% Solvent B (7.5 min), 6-25% B (82.5 min), 25-40% B (30 min), 40-98% B (1 min), and 98% B (15 min). MS1 spectra were acquired in the Orbitrap at 120K resolution with a scan range from 350-1800 m/z, an AGC target of 1e6, and a maximum injection time of 50 ms in Profile mode. MS2 scans were acquired in the Orbitrap at 50K resolution in Centroid mode with the first mass fixed at 110. The cycle time was set to 3 seconds.

LCMS proteomic data were mapped in Scaffold utilizing the Sequest HT search algorithm with the mouse proteome (UniProt UP000000589; 55,485 proteins covering 21,989 genes with a BUSCO assessment of 99.8% genetic coverage). Search parameters were as follows: fully tryptic protease rules with 2 allowed missed cleavages, precursor mass tolerance set to 20 ppm, fragment mass tolerance set to 0.05 Da with only *b* and *y* ions accounted for. Percolator was used as the validation method, based on q-value, with a maximum FDR set to 0.05.

Quantitative analysis is based on TMT MS2 reporter ions generated from HCD fragmentation, with an average reporter S/N threshold of 10, used a co-isolation threshold of 50 with SPS mass matches set at 65%. Normalization was performed at the peptide level, and protein ratios were calculated from the grouped ion abundances, with protein FDR set to a maximum of 0.05.

The mass spectrometry proteomics data are deposited to the ProteomeXchange Consortium via the PRIDE partner repository(*114*). Information on accessing will be made available upon publication of this work.

### Network Medicine Analyses

To start our proteomics analyses, we performed an unbiased Kyoto Encyclopedia of Genes and Genomes (KEGG) Pathway Analysis using ROntoTools R Code,(*49*) similar to our previous study.(*35*) ROnToTools uses overrepresentation analyses as well as functional class scoring to generate significant pathway statistics. This tool was found to have the highest likelihood to identify pathways that were actually changing and the least likely to mis-identify pathways that were not actually changing in a recent systematic review.(*115*) The code was used as is with a modification of ‘hsa’ to ‘mmu’ to reference mouse pathways. Within the ROnToTool algorithm, the Boolean value of total bootstrap permutations accumulation (pAcc) and the Boolean value of over-representation p-value (pORA) were set to 0.01 to determine statistical significance of pathways, as is standard with this code.

To evaluate the temporal effects of pathway dynamics, Temporal TRAnscriptional regulation ModellER (TETRAMER(*116*)) was used to simulate time-based changes in group-based effects. TETRAMER is a publicly available application within Cytoscape that reconstructs gene regulatory networks (GRN) by integrating input temporal data and publicly available GRN and has been used in several previous publications.(*117–120*) Here, we used our mass spectrometry proteomics as input for the temporal component (**Table S3**). We used the Mouse Integrated Protein–Protein Interaction rEference (MIPPIE)(*121*) and a mouse PPI atlas(*122*) as our starting “GRN”. TETRAMER workflow was run exactly as presented in the online tutorial.(*116*) Start and final nodes were determined by the pathways that were found to be significantly changing from our ROnToTools analysis of the VCD animals. For example, since PI3K/Akt Signaling was significantly changing, we input the “starting points” of this pathway, as determined from KEGG pathway as the “start nodes,” and final protein points as the “final nodes” (**Table S2-S3**). We also input the receptors for sex hormones that are likely to be changing with menopause and may be mediating effects. Upregulated proteins were defined as a log2(VCD/SO) > 1, and downregulated proteins were defined as log2(VCD/SO) < -1, which is standard in this analysis.

To verify that observed findings held under a protein-protein interaction network for humans, Protein Interaction Network Analysis For Multiple Sets (PINA4MS) was used to visualize the generated network.(*55, 123*) PINA4MS uses the Human Protein Atlas(*124*) to combine protein-protein interactions and kinase-substrate relationships, and this application was used as presented in the online protocol.

To summarize findings into biological processes, we input the proteins list from Cytoscape in rank order (i.e., from highest fold change to lowest fold change) into g:Profiler.(*57*) To control for the number of proteins within a pathway, the term size was set to a minimum of 5 and a maximum of 5,000. The top 10 biological processes associated with each of the presented networks are displayed in **Table S4**.

Lastly, to simulate the effects of “treating” cartilage with specific sex hormones, we utilized NeDRex to examine changes in proteomic pathways and components. NeDRex is a network-based integrated platform to identify opportunities for drug repurposing.(*66*) Here, the input network was the PPI generated from our TETRAMER simulations. Associations were set to be based off “Pathway-Protein”, “Protein-Protein”, “Drug-Protein”, and “Disorder-Disorder” interactions. Protein-Protein options were set to include self-loop and all evidence. Drug options were listed as “Approved” only, and TrustRank algorithms were used to prioritize here. Specifically, we simulated the interactions of Cetrorelix (LH-receptor antagonist(*67*)), Suramin (FSH-receptor antagonist(*68*)), Serelaxin (relaxin supplement (*69*)), Estradiol, Micronized Progesterone, Levonorgestrel/Estradiol (combined progesterone + estradiol treatment), Dasatinib (senolytic drug (*125*)), and Raloxifene (Selective Estrogen Receptor Modulator).

To quantify node topology of the sex hormone receptors as well as network topology under the different treatment conditions, we used CentiScaPe (*59*). Specifically, we quantified effect size, betweenness, and normalized centroid of the receptors, and diameter, radiality, and closeness of the networks. Procedures for calculating these values was followed as described previously,(*59*) and variable definitions and ranges are listed in **Table S5**.

### Statistical analysis

To determine sample size for histological experiments, an *a priori* power analysis was performed using cartilage histological scoring from previous studies (G*Power).(*126*) For a statistical power of 0.80, a sample size of 5 mice per group was needed. Previous behavioral studies have suggested that 7-9 animals per group are needed to achieve adequate statistical power for the behavioral assays used here.(*127*) Lastly, for the mass spectrometry proteomics study, with a sample size of 5, a statistical power of 90% is maintained for two-fold or greater changes in individual proteins between groups.(*128*)

All statistical analyses were performed in SPSS Statistics for Windows, Version 28.0 (IBM Corp., NY, USA). Statistical significance was defined as an alpha level of 0.05 *a priori*. **Table S7** outlines the statistical tests used for each individual assessment. Where appropriate, normality was assessed using Shapiro-Wilk tests, sphericity was assessed via Mauchly’s test, and equal variance was assessed via Levene’s test.

### Rigor and reproducibility

This study was conducted in accordance with the ARRIVE Guidelines 2.0 (**Supplementary File 3**). In addition to the items mentioned throughout the methods, the methodological protocols were registered *a priori* at The Animal Study Registry (DOI: 10.17590/asr.0000272).(*129*)

## Acknowledgements

The authors would like to thank Dr. Sara Andux, for her excellent veterinary care, and Dr. Lora Rigatti for serving as the blinded pathologist in this study. The behavioral data presented here were collected in the Preclinical Phenotyping Core of the University of Pittsburgh School of Medicine and performed by Gabrielle Little. The mass spectrometry proteomics experiments were performed at the Washington University Proteomics Shared Resource (WU-PSR). The authors gratefully acknowledge the expert technical assistance of Petra Erdmann-Gilmore, Yiling Mi, and Rose Connors as well as Dr. Reid Townsend. The WU-PSR is supported in part by the WU Institute of Clinical and Translational Sciences (NCATS UL1 TR000448), the Mass Spectrometry Research Resource (NIGMS P41 GM103422) and the Siteman Comprehensive Cancer Center Support Grant (NCI P30 CA091842).

## Author contributions

All authors made substantial contributions in the following areas: (1) conception and design of the study, acquisition of data, analysis and interpretation of data, drafting of the article; (2) final approval of the article version to be submitted; and (3) agreement to be personally accountable for the author’s own contributions and to ensure that questions related to the accuracy are appropriately investigated, resolved, and the resolution documented in the literature.

The specific contributions of the authors are as follows:

Conceptualization: GG, HI, FA

Methodology: GG, HI, ZH

Software: GG

Validation: GG, HI, NJ, ACB

Formal Analysis: GG

Investigation: GG, NJ, ACB, JB

Visualization: GG

Funding acquisition: GG, FA

Project administration: GG, FA

Supervision: FA

Writing – original draft: GG

Writing – review & editing: GG, HI, NJ, ZH, ACB, JB, CE, RCT, FA

## Data and materials availability

All data will be made available once the article is published in a peer-reviewed journal.

## Supplementary Figures

**Figure S1:**
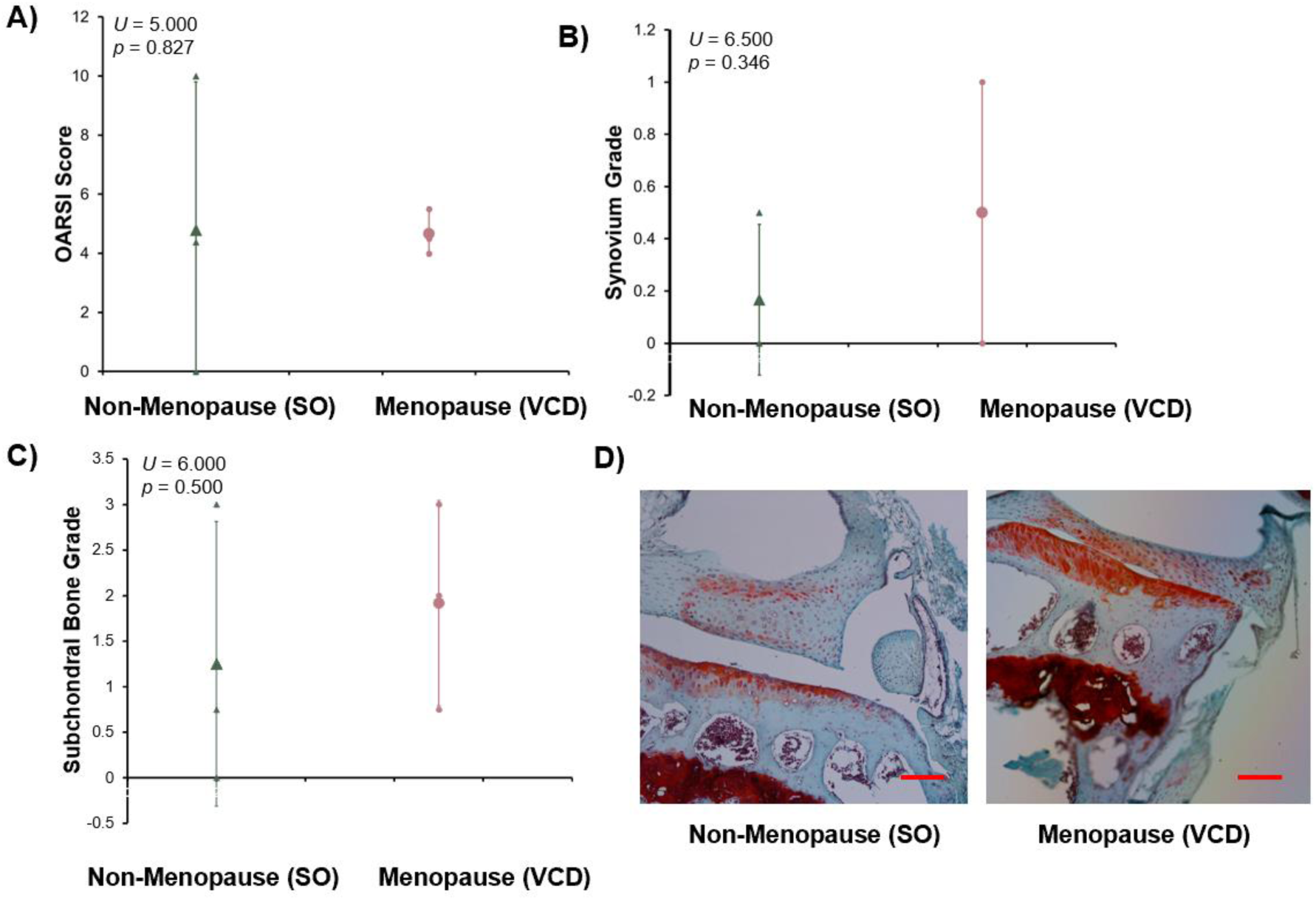
There were no signs of direct effects of sesame oil (SO) and 4-vinylcyclohexene diepoxide (VCD) on the joint tissues. A) Osteoarthritis Research International Society (OARSI) scoring of cartilage degeneration. B) Synovium grade. C) Subchondral bone grade. D) Representative images. Scale bar = 100 μm. Data are presented as mean ± standard deviation. All reagents/materials needed to perform these experiments are listed in **Table S6**, and all statistical tests are listed in **Table S7**.

**Figure S2:**
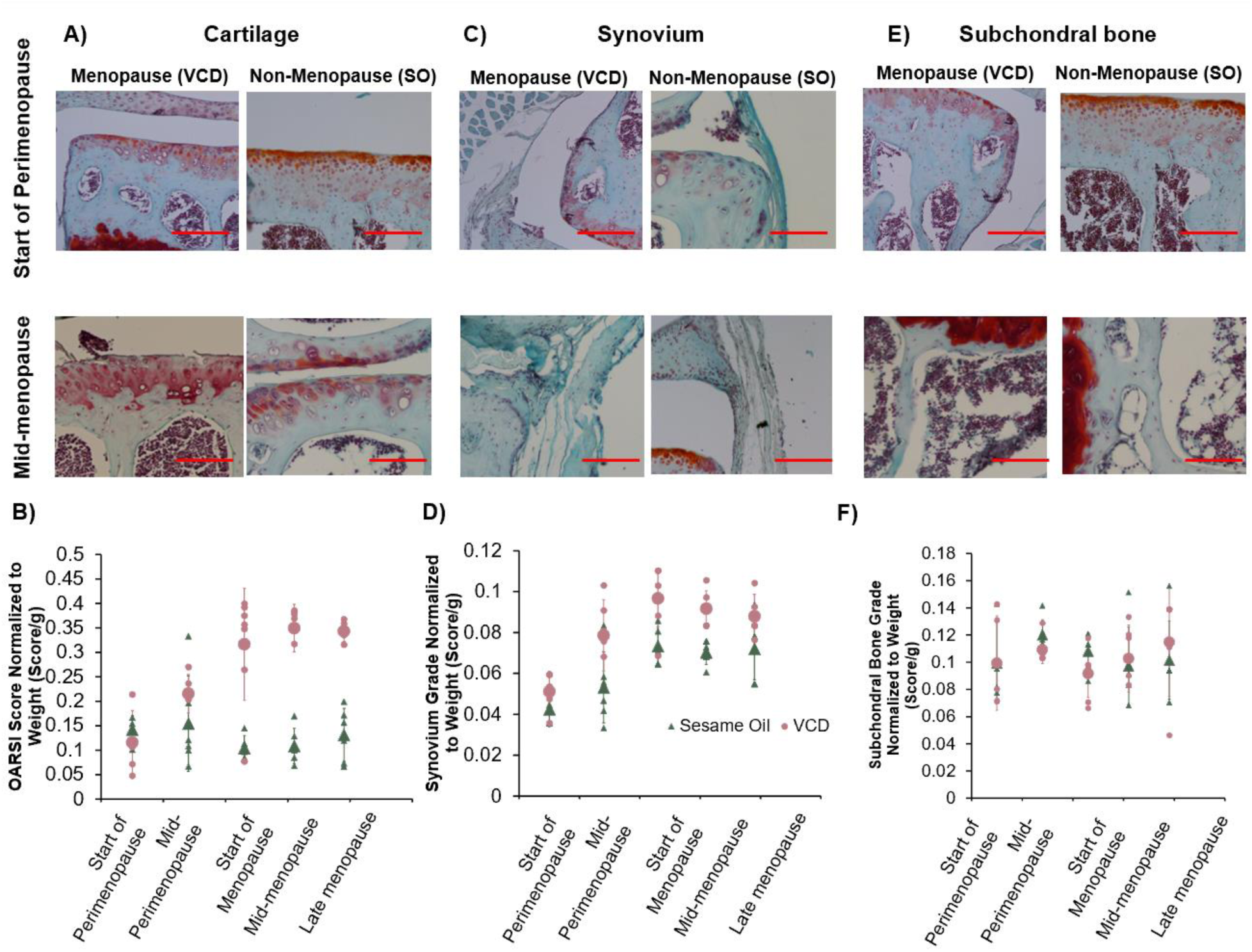
Representative images from menopause and non-menopause groups at the start of perimenopause and mid-menopause and weight normalized scores across groups. A) Cartilage images from start of perimenopause and mid-menopause. B) OARSI score normalized to weight. C) Synovium images from start of perimenopause and mid-menopause. D) Synovium grade normalized to weight. E) Subchondral bone images from start of perimenopause and mid-menopause. F) Subchondral bone grade normalized to weight.

**Figure S3:**
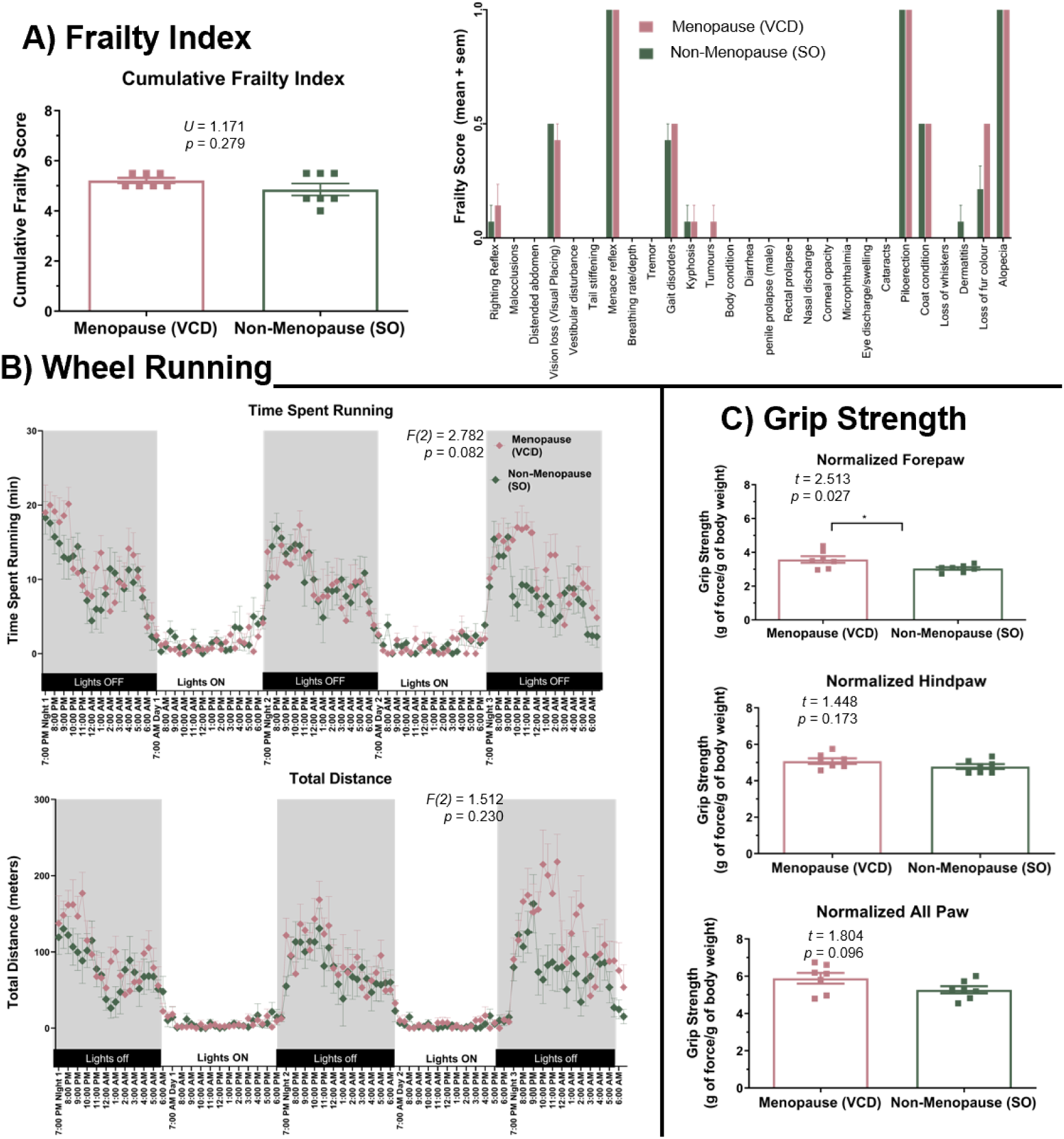
Baseline behavioral characteristics of the groups before treatment with VCD or sesame oil. All behavioral tasks were evaluated prior to starting injections, and data are presented as mean ± standard error of the mean. All reagents/materials needed to perform these experiments are listed in **Table S6**, and all statistical tests are listed in **Table S7**. A) Cumulative frailty index score and individual scores of the different contributing components. Example frailty index scoring sheet is available in **Supplementary File 2**. There was no baseline difference in frailty between groups. B) Time spent and distance run during the wheel running assay. There was no baseline differences in wheel running between groups. C) Forepaw, hindpaw, and all paw grip strength normalized to body weight. There was no baseline difference in hindpaw and all paw grip strength between groups, but forepaw strength was slightly higher in the menopause group compared to the non-menopause group.

**Figure S4:**
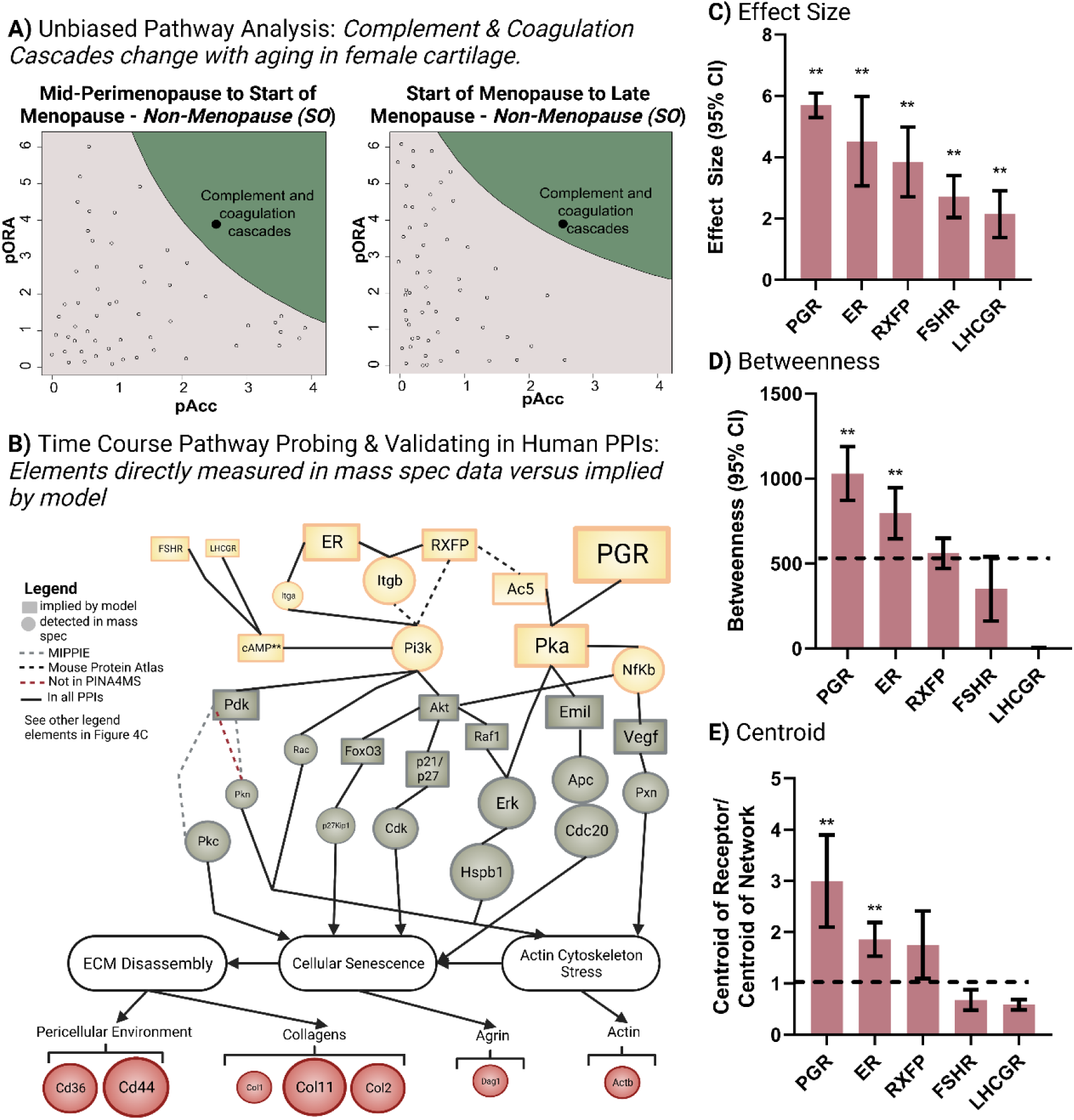
Additional mass spectrometry proteomics results from the unbiased pathway analysis, time course pathway probing, and node topology quantification. A) ROnToTools KEGG Pathway Analysis results from the non-menopause group. B) TETRAMER summary network with only data that were directly measured in mass spectrometry proteomics and demonstrating source of protein-protein interaction (PPI). C) Effect size of progesterone receptor (PGR), estradiol receptor (ER), relaxin receptor (RXFP), follicle stimulating hormone receptor (FSHR), and luteinizing hormone receptor (LHCGR). D) Betweenness (i.e., the capacity of the protein to effectively communicate with distant proteins) of the aforementioned receptors. Dashed line represents betweenness of overall network. E) Centroid (i.e., probability of a protein to organize discrete clusters of proteins) of the aforementioned receptors. Note, here we normalized to the network centroid, and dashed line represents centroid of overall network. ** means p < 0.05. Table S7 lists statistical tests used here. Created in BioRender.

**Figure S5:**
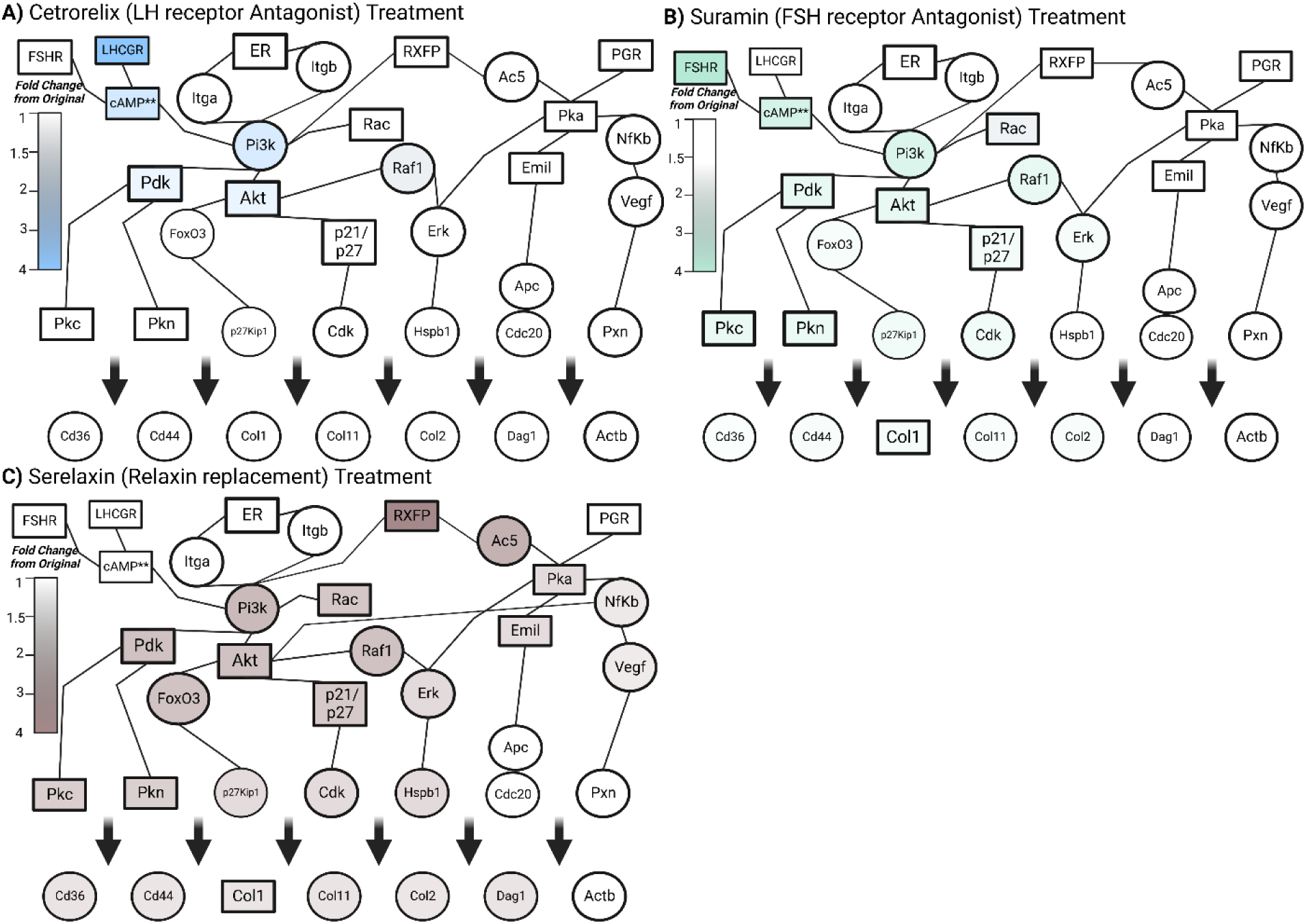
Simulated effects of cetrorelix, suramin, and serelaxin treatment on proteomic network. Fold change is relative to the untreated network (Figure 4C). A) Cetrorelix treatment. B) Suramin treatment. C) Serelaxin treatment. Created in BioRender.

**Figure S6:**
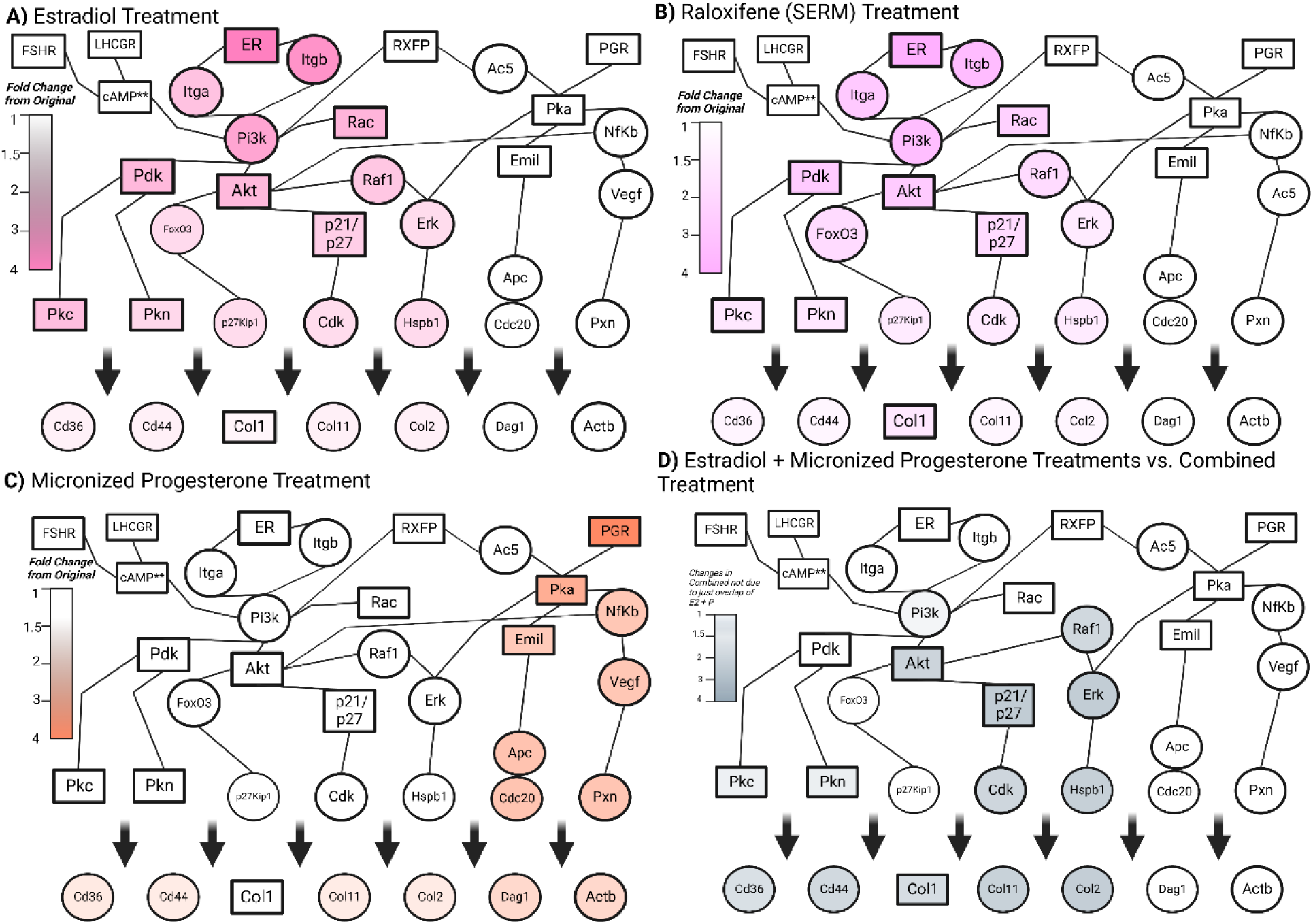
Simulated effects of estradiol, raloxifene, and progesterone treatment on proteomic network. Fold change is relative to the untreated network (Figure 4C). A) Estradiol treatment. B) Raloxifene treatment. C) Micronized Progesterone. D) Difference between combined treatment network and simply adding isolated estradiol and progesterone simulations. Created in BioRender.

## References

1. S. d. Beauvoir, Ralph Ellison Collection (Library of Congress), The second sex. (Knopf, New York,, ed. 1st American, 1953), pp. xxx, 732, xiv p.

2. R. Voskuhl, Sex differences in autoimmune diseases. Biol Sex Differ 2, 1 (2011).

3. N. Reza, J. Gruen, B. Bozkurt, Representation of women in heart failure clinical trials: Barriers to enrollment and strategies to close the gap. Am Heart J Plus 13, (2022).

4. J. Lindenfeld, H. Krause-Steinrauf, J. Salerno, Where are all the women with heart failure? J Am Coll Cardiol 30, 1417–1419 (1997).

5. A. A. Mirin, Gender Disparity in the Funding of Diseases by the U.S. National Institutes of Health. J Womens Health (Larchmt) 30, 956–963 (2021).

6. S. V. Koebele, H. A. Bimonte-Nelson, Modeling menopause: The utility of rodents in translational behavioral endocrinology research. Maturitas 87, 5–17 (2016).

7. R. Diaz Brinton, Minireview: translational animal models of human menopause: challenges and emerging opportunities. Endocrinology 153, 3571–3578 (2012).

8. H. L. Brooks, D. P. Pollow, P. B. Hoyer, The VCD Mouse Model of Menopause and Perimenopause for the Study of Sex Differences in Cardiovascular Disease and the Metabolic Syndrome. Physiology (Bethesda) 31, 250–257 (2016).

9. S. Mohammadalizadeh Charandabi, N. Rezaei, S. Hakimi, A. Montazeri, S. Taheri, H. Taghinejad, K. Sayehmiri, Quality of life of postmenopausal women and their spouses: a community-based study. Iran Red Crescent Med J 17, e21599 (2015).

10. P. The Hormone Therapy Position Statement of The North American Menopause Society” Advisory, The 2022 hormone therapy position statement of The North American Menopause Society. Menopause 29, 767–794 (2022).

11. S. L. Hame, R. A. Alexander, Knee osteoarthritis in women. Curr Rev Musculoskelet Med 6, 182–187 (2013).

12. V. K. Srikanth, J. L. Fryer, G. Zhai, T. M. Winzenberg, D. Hosmer, G. Jones, A meta-analysis of sex differences prevalence, incidence and severity of osteoarthritis. Osteoarthritis Cartilage 13, 769–781 (2005).

13. H. Iijima, G. Gilmer, K. Wang, S. Sivakumar, C. Evans, Y. Matsui, F. Ambrosio, Meta-analysis Integrated With Multi-omics Data Analysis to Elucidate Pathogenic Mechanisms of Age-Related Knee Osteoarthritis in Mice. J Gerontol A Biol Sci Med Sci 77, 1321–1334 (2022).

14. G. Gilmer, A. C. Bean, H. Iijima, N. Jackson, R. C. Thurston, F. Ambrosio, Uncovering the “riddle of femininity” in osteoarthritis: a systematic review and meta-analysis of menopausal animal models and mathematical modeling of estrogen treatment. Osteoarthritis Cartilage, (2023).

15. N. Santoro, Perimenopause: From Research to Practice. J Womens Health (Larchmt) 25, 332–339 (2016).

16. L. T. Shuster, B. S. Gostout, B. R. Grossardt, W. A. Rocca, Prophylactic oophorectomy in premenopausal women and long-term health. Menopause Int 14, 111–116 (2008).

17. D. T. Felson, T. Neogi, Emerging Treatment Models in Rheumatology: Challenges for Osteoarthritis Trials. Arthritis Rheumatol 70, 1175–1181 (2018).

18. D. Chen, J. Shen, W. Zhao, T. Wang, L. Han, J. L. Hamilton, H. J. Im, Osteoarthritis: toward a comprehensive understanding of pathological mechanism. Bone Res 5, 16044 (2017).

19. A. Singh, S. Kaur, I. Walia, A historical perspective on menopause and menopausal age. Bull Indian Inst Hist Med Hyderabad 32, 121–135 (2002).

20. L. E. Wright, P. J. Christian, Z. Rivera, W. G. Van Alstine, J. L. Funk, M. L. Bouxsein, P. B. Hoyer, Comparison of skeletal effects of ovariectomy versus chemically induced ovarian failure in mice. J Bone Miner Res 23, 1296–1303 (2008).

21. F. S. Muhammad, A. K. Goode, N. D. Kock, E. A. Arifin, J. M. Cline, M. R. Adams, P. B. Hoyer, P. J. Christian, S. Isom, J. R. Kaplan, S. E. Appt, Effects of 4-vinylcyclohexene diepoxide on peripubertal and adult Sprague-Dawley rats: ovarian, clinical, and pathologic outcomes. Comp Med 59, 46–59 (2009).

22. A. C. McLean, N. Valenzuela, S. Fai, S. A. Bennett, Performing vaginal lavage, crystal violet staining, and vaginal cytological evaluation for mouse estrous cycle staging identification. J Vis Exp, e4389 (2012).

23. C. S. Caligioni, Assessing reproductive status/stages in mice. Curr Protoc Neurosci Appendix 4, Appendix 4I (2009).

24. L. M. Neff, M. E. Hoffmann, D. M. Zeiss, K. Lowry, M. Edwards, S. M. Rodriguez, K. N. Wachsberg, R. Kushner, L. Landsberg, Core body temperature is lower in postmenopausal women than premenopausal women: potential implications for energy metabolism and midlife weight gain. Cardiovasc Endocrinol 5, 151–154 (2016).

25. G. W. Molnar, Body temperatures during menopausal hot flashes. J Appl Physiol 38, 499–503 (1975).

26. R. R. Freedman, S. Woodward, Core body temperature during menopausal hot flushes. Fertil Steril 65, 1141–1144 (1996).

27. T. Kobayashi, M. Tamura, M. Hayashi, Y. Katsuura, H. Tanabe, T. Ohta, K. Komoriya, Elevation of tail skin temperature in ovariectomized rats in relation to menopausal hot flushes. Am J Physiol Regul Integr Comp Physiol 278, R863–869 (2000).

28. J. M. MacLeay, E. Lehmer, R. M. Enns, C. Mallinckrodt, H. U. Bryant, A. S. Turner, Central and peripheral temperature changes in sheep following ovariectomy. Maturitas 46, 231–238 (2003).

29. R. R. Freedman, D. Norton, S. Woodward, G. Cornelissen, Core body temperature and circadian rhythm of hot flashes in menopausal women. J Clin Endocrinol Metab 80, 2354–2358 (1995).

30. S. Dutta, P. Sengupta, Men and mice: Relating their ages. Life Sci 152, 244–248 (2016).

31. B. Sternfeld, H. Wang, C. P. Quesenberry, Jr., B. Abrams, S. A. Everson-Rose, G. A. Greendale, K. A. Matthews, J. I. Torrens, M. Sowers, Physical activity and changes in weight and waist circumference in midlife women: findings from the Study of Women’s Health Across the Nation. Am J Epidemiol 160, 912–922 (2004).

32. C. Karvonen-Gutierrez, C. Kim, Association of Mid-Life Changes in Body Size, Body Composition and Obesity Status with the Menopausal Transition. Healthcare (Basel) 4, (2016).

33. A. Fenton, Weight, Shape, and Body Composition Changes at Menopause. J Midlife Health 12, 187–192 (2021).

34. V. B. Kraus, J. L. Huebner, J. DeGroot, A. Bendele, The OARSI histopathology initiative - recommendations for histological assessments of osteoarthritis in the guinea pig. Osteoarthritis Cartilage 18 Suppl 3, S35–52 (2010).

35. H. Iijima, G. Gilmer, K. Wang, A. C. Bean, Y. He, H. Lin, W. Y. Tang, D. Lamont, C. Tai, A. Ito, J. J. Jones, C. Evans, F. Ambrosio, Age-related matrix stiffening epigenetically regulates alpha-Klotho expression and compromises chondrocyte integrity. Nat Commun 14, 18 (2023).

36. K. M. Leyland, D. J. Hart, M. K. Javaid, A. Judge, A. Kiran, A. Soni, L. M. Goulston, C. Cooper, T. D. Spector, N. K. Arden, The natural history of radiographic knee osteoarthritis: a fourteen-year population-based cohort study. Arthritis Rheum 64, 2243–2251 (2012).

37. D. T. Felson, Y. Zhang, M. T. Hannan, A. Naimark, B. N. Weissman, P. Aliabadi, D. Levy, The incidence and natural history of knee osteoarthritis in the elderly. The Framingham Osteoarthritis Study. Arthritis Rheum 38, 1500–1505 (1995).

38. J. Cibere, E. C. Sayre, A. Guermazi, S. Nicolaou, J. A. Kopec, J. M. Esdaile, A. Thorne, J. Singer, H. Wong, Natural history of cartilage damage and osteoarthritis progression on magnetic resonance imaging in a population-based cohort with knee pain. Osteoarthritis Cartilage 19, 683–688 (2011).

39. L. Lachance, M. F. Sowers, D. Jamadar, M. Hochberg, The natural history of emergent osteoarthritis of the knee in women. Osteoarthritis Cartilage 10, 849–854 (2002).

40. V. B. Kraus, J. L. Huebner, J. DeGroot, A. Bendele, The OARSI histopathology initiative - recommendations for histological assessments of osteoarthritis in the guinea pig. Osteoarthritis Cartilage 18 Suppl 3, S35–52 (2010).

41. L. B. Cao, C. K. Leung, P. W. Law, Y. Lv, C. H. Ng, H. B. Liu, G. Lu, J. L. Ma, W. Y. Chan, Systemic changes in a mouse model of VCD-induced premature ovarian failure. Life Sci 262, 118543 (2020).

42. X. Xu, X. Li, Y. Liang, Y. Ou, J. Huang, J. Xiong, L. Duan, D. Wang, Estrogen Modulates Cartilage and Subchondral Bone Remodeling in an Ovariectomized Rat Model of Postmenopausal Osteoarthritis. Med Sci Monit 25, 3146–3153 (2019).

43. F. M. Reis, N. Pestana-Oliveira, C. M. Leite, F. B. Lima, M. L. Brandao, F. G. Graeff, C. M. Del- Ben, J. A. Anselmo-Franci, Hormonal changes and increased anxiety-like behavior in a perimenopause-animal model induced by 4-vinylcyclohexene diepoxide (VCD) in female rats. Psychoneuroendocrinology 49, 130–140 (2014).

44. D. D. Dunlop, J. Song, P. A. Semanik, L. Sharma, R. W. Chang, Physical activity levels and functional performance in the osteoarthritis initiative: a graded relationship. Arthritis Rheum 63, 127–136 (2011).

45. C. Gay, C. Guiguet-Auclair, C. Mourgues, L. Gerbaud, E. Coudeyre, Physical activity level and association with behavioral factors in knee osteoarthritis. Ann Phys Rehabil Med 62, 14–20 (2019).

46. A. H. Lee, K. B. Detweiler, T. A. Harper, K. E. Knap, M. R. C. de Godoy, K. S. Swanson, Physical activity patterns of free living dogs diagnosed with osteoarthritis. J Anim Sci 99, (2021).

47. M. Gosset, F. Berenbaum, S. Thirion, C. Jacques, Primary culture and phenotyping of murine chondrocytes. Nat Protoc 3, 1253–1260 (2008).

48. J. Zhao, Q. Ouyang, Z. Hu, Q. Huang, J. Wu, R. Wang, M. Yang, A protocol for the culture and isolation of murine synovial fibroblasts. Biomed Rep 5, 171–175 (2016).

49. A. S. Voichita C, Draghici S, R. p. version, Ed. (2022).

50. K. Bubb, T. Holzer, J. L. Nolte, M. Kruger, R. Wilson, U. Schlotzer-Schrehardt, J. Brinckmann, J. Altmuller, A. Aszodi, L. Fleischhauer, H. Clausen-Schaumann, K. Probst, B. Brachvogel, Mitochondrial respiratory chain function promotes extracellular matrix integrity in cartilage. J Biol Chem 297, 101224 (2021).

51. A. Naba, O. M. T. Pearce, A. Del Rosario, D. Ma, H. Ding, V. Rajeeve, P. R. Cutillas, F. R. Balkwill, R. O. Hynes, Characterization of the Extracellular Matrix of Normal and Diseased Tissues Using Proteomics. J Proteome Res 16, 3083–3091 (2017).

52. N. Santoro, J. R. Brown, T. Adel, J. H. Skurnick, Characterization of reproductive hormonal dynamics in the perimenopause. J Clin Endocrinol Metab 81, 1495–1501 (1996).

53. A. L. Barabasi, N. Gulbahce, J. Loscalzo, Network medicine: a network-based approach to human disease. Nat Rev Genet 12, 56–68 (2011).

54. P. E. Cholley, J. Moehlin, A. Rohmer, V. Zilliox, S. Nicaise, H. Gronemeyer, M. A. Mendoza- Parra, Modeling gene-regulatory networks to describe cell fate transitions and predict master regulators. NPJ Syst Biol Appl 4, 29 (2018).

55. J. Wu, T. Vallenius, K. Ovaska, J. Westermarck, T. P. Makela, S. Hautaniemi, Integrated network analysis platform for protein-protein interactions. Nat Methods 6, 75–77 (2009).

56. M. J. Cowley, M. Pinese, K. S. Kassahn, N. Waddell, J. V. Pearson, S. M. Grimmond, A. V. Biankin, S. Hautaniemi, J. Wu, PINA v2.0: mining interactome modules. Nucleic Acids Res 40, D862–865 (2012).

57. U. Raudvere, L. Kolberg, I. Kuzmin, T. Arak, P. Adler, H. Peterson, J. Vilo, g:Profiler: a web server for functional enrichment analysis and conversions of gene lists (2019 update). Nucleic Acids Res 47, W191–W198 (2019).

58. J. Reimand, R. Isserlin, V. Voisin, M. Kucera, C. Tannus-Lopes, A. Rostamianfar, L. Wadi, M. Meyer, J. Wong, C. Xu, D. Merico, G. D. Bader, Pathway enrichment analysis and visualization of omics data using g:Profiler, GSEA, Cytoscape and EnrichmentMap. Nat Protoc 14, 482–517 (2019).

59. G. Scardoni, M. Petterlini, C. Laudanna, Analyzing biological network parameters with CentiScaPe. Bioinformatics 25, 2857–2859 (2009).

60. J. A. Roman-Blas, S. Castaneda, R. Largo, G. Herrero-Beaumont, Osteoarthritis associated with estrogen deficiency. Arthritis Res Ther 11, 241 (2009).

61. Y. H. Sniekers, H. Weinans, G. J. van Osch, J. P. van Leeuwen, Oestrogen is important for maintenance of cartilage and subchondral bone in a murine model of knee osteoarthritis. Arthritis Res Ther 12, R182 (2010).

62. R. Dreier, T. Ising, M. Ramroth, Y. Rellmann, Estradiol Inhibits ER Stress-Induced Apoptosis in Chondrocytes and Contributes to a Reduced Osteoarthritic Cartilage Degeneration in Female Mice. Front Cell Dev Biol 10, 913118 (2022).

63. Y. Liu, M. Zhang, D. Kong, Y. Wang, J. Li, W. Liu, Y. Fu, J. Xu, High follicle-stimulating hormone levels accelerate cartilage damage of knee osteoarthritis in postmenopausal women through the PI3K/AKT/NF-kappaB pathway. FEBS Open Bio 10, 2235–2245 (2020).

64. Z. Huan, Y. Wang, M. Zhang, X. Zhang, Y. Liu, L. Kong, J. Xu, Follicle-stimulating hormone worsens osteoarthritis by causing inflammation and chondrocyte dedifferentiation. FEBS Open Bio 11, 2292–2303 (2021).

65. Y. Wang, M. Zhang, Z. Huan, S. Shao, X. Zhang, D. Kong, J. Xu, FSH directly regulates chondrocyte dedifferentiation and cartilage development. J Endocrinol 248, 193–206 (2021).

66. S. Sadegh, J. Skelton, E. Anastasi, J. Bernett, D. B. Blumenthal, G. Galindez, M. Salgado- Albarran, O. Lazareva, K. Flanagan, S. Cockell, C. Nogales, A. I. Casas, H. Schmidt, J. Baumbach, A. Wipat, T. Kacprowski, Network medicine for disease module identification and drug repurposing with the NeDRex platform. Nat Commun 12, 6848 (2021).

67. P. van Gastel, M. van der Zanden, D. Telting, M. Filius, L. Bancsi, H. de Boer, Luteinizing hormone-releasing hormone receptor antagonist may reduce postmenopausal flushing. Menopause 19, 178–185 (2012).

68. R. C. Anderson, C. L. Newton, R. P. Millar, Small Molecule Follicle-Stimulating Hormone Receptor Agonists and Antagonists. Front Endocrinol (Lausanne) 9, 757 (2018).

69. C. S. Samuel, T. D. Hewitson, E. N. Unemori, M. L. Tang, Drugs of the future: the hormone relaxin. Cell Mol Life Sci 64, 1539–1557 (2007).

70. J. L. Kirkland, T. Tchkonia, Senolytic drugs: from discovery to translation. J Intern Med 288, 518–536 (2020).

71. A. Mathiessen, P. G. Conaghan, Synovitis in osteoarthritis: current understanding with therapeutic implications. Arthritis Res Ther 19, 18 (2017).

72. Z. Wu, S. H. Korntner, A. M. Mullen, D. I. Zeugolis, Collagen type II: From biosynthesis to advanced biomaterials for cartilage engineering. Biomaterials and Biosystems 4, 100030 (2021).

73. E. V. Raine, A. W. Dodd, L. N. Reynard, J. Loughlin, Allelic expression analysis of the osteoarthritis susceptibility gene COL11A1 in human joint tissues. BMC Musculoskelet Disord 14, 85 (2013).

74. J. C. Sok, J. A. Lee, S. Dasari, S. Joyce, S. C. Contrucci, A. M. Egloff, B. K. Trevelline, R. Joshi, N. Kumari, J. R. Grandis, S. M. Thomas, Collagen type XI alpha1 facilitates head and neck squamous cell cancer growth and invasion. Br J Cancer 109, 3049–3056 (2013).

75. V. W. Tang, Collagen, stiffness, and adhesion: the evolutionary basis of vertebrate mechanobiology. Mol Biol Cell 31, 1823–1834 (2020).

76. S. M. Smith, D. E. Birk, Focus on molecules: collagens V and XI. Exp Eye Res 98, 105–106 (2012).

77. Y. Gao, S. Liu, J. Huang, W. Guo, J. Chen, L. Zhang, B. Zhao, J. Peng, A. Wang, Y. Wang, W. Xu, S. Lu, M. Yuan, Q. Guo, The ECM-cell interaction of cartilage extracellular matrix on chondrocytes. Biomed Res Int 2014, 648459 (2014).

78. K. Sun, J. Luo, J. Guo, X. Yao, X. Jing, F. Guo, The PI3K/AKT/mTOR signaling pathway in osteoarthritis: a narrative review. Osteoarthritis Cartilage 28, 400–409 (2020).

79. A. Mobasheri, M. P. Rayman, O. Gualillo, J. Sellam, P. van der Kraan, U. Fearon, The role of metabolism in the pathogenesis of osteoarthritis. Nat Rev Rheumatol 13, 302–311 (2017).

80. S. C. Rosa, A. T. Rufino, F. Judas, C. Tenreiro, M. C. Lopes, A. F. Mendes, Expression and function of the insulin receptor in normal and osteoarthritic human chondrocytes: modulation of anabolic gene expression, glucose transport and GLUT-1 content by insulin. Osteoarthritis Cartilage 19, 719–727 (2011).

81. E. M. Ciruelos Gil, Targeting the PI3K/AKT/mTOR pathway in estrogen receptor-positive breast cancer. Cancer Treat Rev 40, 862–871 (2014).

82. G. Boccalini, C. Sassoli, D. Bani, S. Nistri, Relaxin induces up-regulation of ADAM10 metalloprotease in RXFP1-expressing cells by PI3K/AKT signaling. Mol Cell Endocrinol 472, 80–86 (2018).

83. A. L. Valkovic, R. A. Bathgate, C. S. Samuel, M. Kocan, Understanding relaxin signalling at the cellular level. Mol Cell Endocrinol 487, 24–33 (2019).

84. C. Schwabe, E. E. Bullesbach, Relaxin. Comp Biochem Physiol B 96, 15–21 (1990).

85. J. Bonaventure, B. de La Tour, L. Tsagris, L. W. Eddie, G. Tregear, M. T. Corvol, Effect of relaxin on the phenotype of collagens synthesized by cultured rabbit chondrocytes. Biochim Biophys Acta 972, 209–220 (1988).

86. F. Dehghan, B. S. Haerian, S. Muniandy, A. Yusof, J. L. Dragoo, N. Salleh, The effect of relaxin on the musculoskeletal system. Scand J Med Sci Sports 24, e220–229 (2014).

87. M. P. Hellio Le Graverand, C. Reno, D. A. Hart, Influence of pregnancy on gene expression in rabbit articular cartilage. Osteoarthritis Cartilage 6, 341–350 (1998).

88. L. S. Trevino, W. E. Bingman, 3rd, D. P. Edwards, W. Nl, The requirement for p42/p44 MAPK activity in progesterone receptor-mediated gene regulation is target gene-specific. Steroids 78, 542–547 (2013).

89. M. Salazar, A. Lerma-Ortiz, G. M. Hooks, A. K. Ashley, R. L. Ashley, Progestin-mediated activation of MAPK and AKT in nuclear progesterone receptor negative breast epithelial cells: The role of membrane progesterone receptors. Gene 591, 6–13 (2016).

90. A. R. Dwyer, T. H. Truong, J. H. Ostrander, C. A. Lange, 90 YEARS OF PROGESTERONE: Steroid receptors as MAPK signaling sensors in breast cancer: let the fates decide. J Mol Endocrinol 65, T35–T48 (2020).

91. A. Skildum, E. Faivre, C. A. Lange, Progesterone receptors induce cell cycle progression via activation of mitogen-activated protein kinases. Mol Endocrinol 19, 327–339 (2005).

92. Wardhana, E. E. Surachmanto, E. A. Datau, J. Ongkowijaya, A. M. Karema-Kaparang, Transdermal bio-identical progesterone cream as hormonal treatment for osteoarthritis. Acta Med Indones 45, 224–232 (2013).

93. M. C. Nevitt, D. T. Felson, E. N. Williams, D. Grady, The effect of estrogen plus progestin on knee symptoms and related disability in postmenopausal women: The Heart and Estrogen/Progestin Replacement Study, a randomized, double-blind, placebo-controlled trial. Arthritis Rheum 44, 811–818 (2001).

94. J. Xie, Y. Wang, L. Lu, L. Liu, X. Yu, F. Pei, Cellular senescence in knee osteoarthritis: molecular mechanisms and therapeutic implications. Ageing Res Rev 70, 101413 (2021).

95. R. F. Loeser, Aging and osteoarthritis: the role of chondrocyte senescence and aging changes in the cartilage matrix. Osteoarthritis Cartilage 17, 971–979 (2009).

96. K. McCulloch, G. J. Litherland, T. S. Rai, Cellular senescence in osteoarthritis pathology. Aging Cell 16, 210–218 (2017).

97. O. H. Jeon, N. David, J. Campisi, J. H. Elisseeff, Senescent cells and osteoarthritis: a painful connection. J Clin Invest 128, 1229–1237 (2018).

98. M. L. Ji, H. Jiang, Z. Li, R. Geng, J. Z. Hu, Y. C. Lin, J. Lu, Sirt6 attenuates chondrocyte senescence and osteoarthritis progression. Nat Commun 13, 7658 (2022).

99. Y. Liu, Z. Zhang, T. Li, H. Xu, H. Zhang, Senescence in osteoarthritis: from mechanism to potential treatment. Arthritis Res Ther 24, 174 (2022).

100. Q. Guo, X. Chen, J. Chen, G. Zheng, C. Xie, H. Wu, Z. Miao, Y. Lin, X. Wang, W. Gao, X. Zheng, Z. Pan, Y. Zhou, Y. Wu, X. Zhang, STING promotes senescence, apoptosis, and extracellular matrix degradation in osteoarthritis via the NF-kappaB signaling pathway. Cell Death Dis 12, 13 (2021).

101. L. Deng, R. Ren, Z. Liu, M. Song, J. Li, Z. Wu, X. Ren, L. Fu, W. Li, W. Zhang, P. Guillen, J. C. Izpisua Belmonte, P. Chan, J. Qu, G. H. Liu, Stabilizing heterochromatin by DGCR8 alleviates senescence and osteoarthritis. Nat Commun 10, 3329 (2019).

102. T. C. Beadnell, K. W. Nassar, M. M. Rose, E. G. Clark, B. P. Danysh, M. C. Hofmann, N. Pozdeyev, R. E. Schweppe, Src-mediated regulation of the PI3K pathway in advanced papillary and anaplastic thyroid cancer. Oncogenesis 7, 23 (2018).

103. J. Lei, D. H. Ingbar, Src kinase integrates PI3K/Akt and MAPK/ERK1/2 pathways in T3-induced Na-K-ATPase activity in adult rat alveolar cells. Am J Physiol Lung Cell Mol Physiol 301, L765–771 (2011).

104. H. S. Lee, C. Moon, H. W. Lee, E. M. Park, M. S. Cho, J. L. Kang, Src tyrosine kinases mediate activations of NF-kappaB and integrin signal during lipopolysaccharide-induced acute lung injury. J Immunol 179, 7001–7011 (2007).

105. A. Migliaccio, D. Piccolo, G. Castoria, M. Di Domenico, A. Bilancio, M. Lombardi, W. Gong, M. Beato, F. Auricchio, Activation of the Src/p21ras/Erk pathway by progesterone receptor via cross-talk with estrogen receptor. EMBO J 17, 2008–2018 (1998).

106. D. J. Hunter, J. J. McDougall, F. J. Keefe, The symptoms of osteoarthritis and the genesis of pain. Rheum Dis Clin North Am 34, 623–643 (2008).

107. A. R. Sonawane, S. T. Weiss, K. Glass, A. Sharma, Network Medicine in the Age of Biomedical Big Data. Front Genet 10, 294 (2019).

108. F. Sensini, D. Inta, R. Palme, C. Brandwein, N. Pfeiffer, M. A. Riva, P. Gass, A. S. Mallien, The impact of handling technique and handling frequency on laboratory mouse welfare is sex-specific. Sci Rep 10, 17281 (2020).

109. D. Hayashi, D. T. Felson, J. Niu, D. J. Hunter, F. W. Roemer, P. Aliabadi, A. Guermazi, Pre-radiographic osteoarthritic changes are highly prevalent in the medial patella and medial posterior femur in older persons: Framingham OA study. Osteoarthritis Cartilage 22, 76–83 (2014).

110. A. R. Armstrong, C. S. Carlson, A. K. Rendahl, R. F. Loeser, Optimization of histologic grading schemes in spontaneous and surgically-induced murine models of osteoarthritis. Osteoarthritis Cartilage 29, 536–546 (2021).

111. S. S. Glasson, M. G. Chambers, W. B. Van Den Berg, C. B. Little, The OARSI histopathology initiative - recommendations for histological assessments of osteoarthritis in the mouse. Osteoarthritis Cartilage 18 Suppl 3, S17–23 (2010).

112. S. J. Sukoff Rizzo, L. C. Anderson, T. L. Green, T. McGarr, G. Wells, S. S. Winter, Assessing Healthspan and Lifespan Measures in Aging Mice: Optimization of Testing Protocols, Replicability, and Rater Reliability. Curr Protoc Mouse Biol 8, e45 (2018).

113. M. D. Gardiner, T. L. Vincent, C. Driscoll, A. Burleigh, G. Bou-Gharios, J. Saklatvala, H. Nagase, A. Chanalaris, Transcriptional analysis of micro-dissected articular cartilage in post-traumatic murine osteoarthritis. Osteoarthritis Cartilage 23, 616–628 (2015).

114. Y. Perez-Riverol, A. Csordas, J. Bai, M. Bernal-Llinares, S. Hewapathirana, D. J. Kundu, A. Inuganti, J. Griss, G. Mayer, M. Eisenacher, E. Pérez, J. Uszkoreit, J. Pfeuffer, T. Sachsenberg, S. Yilmaz, S. Tiwary, J. Cox, E. Audain, M. Walzer, A. F. Jarnuczak, T. Ternent, A. Brazma, J. A. Vizcaíno, The PRIDE database and related tools and resources in 2019: improving support for quantification data. Nucleic Acids Res 47, D442–d450 (2019).

115. T. M. Nguyen, A. Shafi, T. Nguyen, S. Draghici, Identifying significantly impacted pathways: a comprehensive review and assessment. Genome Biol 20, 203 (2019).

116. 116. P.-E. C. Marco-Antonio Mendoza-Parra, Julien Moehlin, Alexia Rohmer, Vincent Zilliox and Hinrich Gronemeyer. (2023).

117. A. C. D’Alessio, Z. P. Fan, K. J. Wert, P. Baranov, M. A. Cohen, J. S. Saini, E. Cohick, C. Charniga, D. Dadon, N. M. Hannett, M. J. Young, S. Temple, R. Jaenisch, T. I. Lee, R. A. Young, A Systematic Approach to Identify Candidate Transcription Factors that Control Cell Identity. Stem Cell Reports 5, 763–775 (2015).

118. M. Koga, M. Matsuda, T. Kawamura, T. Sogo, A. Shigeno, E. Nishida, M. Ebisuya, Foxd1 is a mediator and indicator of the cell reprogramming process. Nat Commun 5, 3197 (2014).

119. F. Rapino, E. F. Robles, J. A. Richter-Larrea, E. M. Kallin, J. A. Martinez-Climent, T. Graf, C/EBPalpha induces highly efficient macrophage transdifferentiation of B lymphoma and leukemia cell lines and impairs their tumorigenicity. Cell Rep 3, 1153–1163 (2013).

120. M. A. Mendoza-Parra, V. Malysheva, M. A. Mohamed Saleem, M. Lieb, A. Godel, H. Gronemeyer, Reconstructed cell fate-regulatory programs in stem cells reveal hierarchies and key factors of neurogenesis. Genome Res 26, 1505–1519 (2016).

121. G. Alanis-Lobato, J. S. Mollmann, M. H. Schaefer, M. A. Andrade-Navarro, MIPPIE: the mouse integrated protein-protein interaction reference. Database (Oxford) 2020, (2020).

122. M. A. Skinnider, N. E. Scott, A. Prudova, C. H. Kerr, N. Stoynov, R. G. Stacey, Q. W. T. Chan, D. Rattray, J. Gsponer, L. J. Foster, An atlas of protein-protein interactions across mouse tissues. Cell 184, 4073–4089 e4017 (2021).

123. Y. Du, M. Cai, X. Xing, J. Ji, E. Yang, J. Wu, PINA 3.0: mining cancer interactome. Nucleic Acids Res 49, D1351–D1357 (2021).

124. M. Uhlen, L. Fagerberg, B. M. Hallstrom, C. Lindskog, P. Oksvold, A. Mardinoglu, A. Sivertsson, C. Kampf, E. Sjostedt, A. Asplund, I. Olsson, K. Edlund, E. Lundberg, S. Navani, C. A. Szigyarto, J. Odeberg, D. Djureinovic, J. O. Takanen, S. Hober, T. Alm, P. H. Edqvist, H. Berling, H. Tegel, J. Mulder, J. Rockberg, P. Nilsson, J. M. Schwenk, M. Hamsten, K. von Feilitzen, M. Forsberg, L. Persson, F. Johansson, M. Zwahlen, G. von Heijne, J. Nielsen, F. Ponten, Proteomics. Tissue-based map of the human proteome. Science 347, 1260419 (2015).

125. T. D. Saccon, R. Nagpal, H. Yadav, M. B. Cavalcante, A. D. C. Nunes, A. Schneider, A. Gesing, B. Hughes, M. Yousefzadeh, T. Tchkonia, J. L. Kirkland, L. J. Niedernhofer, P. D. Robbins, M. M. Masternak, Senolytic Combination of Dasatinib and Quercetin Alleviates Intestinal Senescence and Inflammation and Modulates the Gut Microbiome in Aged Mice. J Gerontol A Biol Sci Med Sci 76, 1895–1905 (2021).

126. P. Hoegh-Andersen, L. B. Tanko, T. L. Andersen, C. V. Lundberg, J. A. Mo, A. M. Heegaard, J. M. Delaisse, S. Christgau, Ovariectomized rats as a model of postmenopausal osteoarthritis: validation and application. Arthritis Res Ther 6, R169–180 (2004).

127. S. J. Sukoff Rizzo, J. L. Silverman, Methodological Considerations for Optimizing and Validating Behavioral Assays. Curr Protoc Mouse Biol 6, 364–379 (2016).

128. Y. Levin, The role of statistical power analysis in quantitative proteomics. Proteomics 11, 2565–2567 (2011).

129. B. Bert, C. Heinl, J. Chmielewska, F. Schwarz, B. Grune, A. Hensel, M. Greiner, G. Schonfelder, Refining animal research: The Animal Study Registry. PLoS Biol 17, e3000463 (2019).

